# Targeted hydrolysis of native potato protein: A novel route for obtaining hydrolysates with improved interfacial properties

**DOI:** 10.1101/2022.05.25.493405

**Authors:** Simon Gregersen Echers, Ali Jafarpour, Betül Yesiltas, Pedro J. García-Moreno, Mathias Greve-Poulsen, Dennis Hansen, Charlotte Jacobsen, Michael Toft Overgaard, Egon Bech Hansen

**Author notes:** Corresponding author (SGE). Simon Gregersen Echers and Ali Jafarpour contributed equally to this work.

## Abstract

Peptides and protein hydrolysates are promising alternatives to substitute chemical additives as functional food ingredients. In this study, we present a novel approach for producing a potato protein hydrolysate with improved emulsifying and foaming properties by data-driven, targeted hydrolysis. Based on previous studies, we selected 15 emulsifier peptides derived from abundant potato proteins, which were clustered based on sequence identity. Through *in silico* analysis, we determined that from a range of industrial proteases (Neutrase (Neut), Alcalase (Alc), Flavorzyme (Flav) and Trypsin (Tryp)), Tryp was found more likely to release peptides resembling the target peptides. After applying all proteases individually, hydrolysates were assayed for *in vitro* emulsifying and foaming properties. No direct correlation between degree of hydrolysis and interfacial properties was found. Tryp produced a hydrolysate (DH=5.4%) with the highest (P<0.05) emulsifying and foaming abilities, good stabilities, and high aqueous solubility. Using LC-MS/MS, we identified >10,000 peptides in each hydrolysate. Through peptide mapping, we show that random overlapping with known peptide emulsifiers is not sufficient to quantitatively describe hydrolysate functionality. While Neut hydrolysates had the highest proportion of peptides with target overlap, they showed inferior interfacial activity. In contrast, Tryp was able to release specifically targeted peptides, explaining the high surface activity observed. While modest yields and residual unhydrolyzed protein indicate room for process improvement, this work shows that data-driven, targeted hydrolysis is a viable, interdisciplinary approach to facilitate hydrolysis design for production of functional hydrolysates from alternative protein sources.

## 1. Introduction

Potato (*Solanum tuberosum*) is the fourth most cultivated crop with a global production of about 370 million tonnes in 2018 (Food and Agriculture Organization of the United Nations, 2020). Potatoes are the second highest protein providing crop per area grown after wheat, and despite a modest protein content of 1-2% depending on cultivar (Camire, Kubow, & Donnelly, 2009; Jørgensen, Stensballe, & Welinder, 2011; Van Koningsveld et al., 2006), they are still regarded as a highly attractive food protein due to both high nutritional quality and functional properties (Waglay & Karboune, 2016b). Potato proteins are often classified according to their cellular function with patatin, also known as tuberin, as the major fraction. Patatins are highly homologous storage proteins with molecular weights (MWs) from 40 to 45 kDa and pIs in the range of 4.8-5.2 (Kärenlampi & White, 2009), which constitute 35-40% of the tuber protein depending on the specific cultivar (Løkra & Strætkvern, 2009). Likewise, protease inhibitors constitute 30–40% of the total tuber protein (Bauw et al., 2006; Jørgensen, Bauw, & Welinder, 2006), but represent a group of more diverse proteins with MWs from 5 to 25 kDa (Heibges, Glaczinski, Ballvora, Salamini, & Gebhardt, 2003; Pouvreau et al., 2001) which can be divided into sub-groups based on sequence homology and thus targets for inhibition (García-Moreno, Gregersen, et al., 2020; Heibges et al., 2003).

Directly isolated from potato fruit juice (PFJ), native potato protein has been reported to exhibit high solubility as well as foaming and emulsifying activity, which has primarily been ascribed to the large patatin content (Ralet & Guéguen, 2000; Schmidt et al., 2019; Van Koningsveld et al., 2001, 2006). To achieve these desirable characteristics, it is important to use appropriate extraction methods to maintain the proteins in their native and intact form, which, from the industrial point of view, would be costly. Thus, the industrially isolated potato protein, mainly obtained in denatured form through rather harsh heat coagulation and acid precipitation, lacks those above-mentioned functionalities. However, due to high content of amino acids with hydrophobic functional groups, in particular, with branched (isoleucine, leucine, and valine) and aromatic (phenylalanine and tyrosine) side chains (Refstie & Tiekstra, 2003), techno-functionality of potato protein can potentially be improved when the large, denatured proteins undergo a specific set of hydrolysis steps to yield smaller peptides (Aluko, 2018; Li-Chan, 2015; Moreno, Cuadrado, Marquez Moreno, & Fernandez Cuadrado, 1993; Rodan, Fields, & Falla, 2013; Wang & Xiong, 2005). Enzymatically released peptides may display better functional properties than their parent protein molecules and consequently exhibit higher activity in food systems (Kamnerdpetch, Weiss, Kasper, & Scheper, 2007; Moreno et al., 1993). Moreover, potato protein hydrolysates have also been shown to have beneficial health effects *in vivo* (Chuang et al., 2020), illustrating their potential as both a functional and bioactive ingredients.

Amphiphilic surfactants are widely used as emulsifiers as they contain both hydrophobic and hydrophilic regions, which are capable of reorganising at the oil-water interface and thereby stabilize the emulsion by decreasing the interfacial tension between the two immiscible liquids (McClements & Jafari, 2018). In this respect, the use of peptides as natural emulsifiers and biosurfactants has received increasing attention over the past few decades from both the academic and the industrial sector (Adjonu, Doran, Torley, & Agboola, 2014; Dexter & Middelberg, 2008; Hanley & James, 2018; Le Guenic, Chaveriat, Lequart, Joly, & Martin, 2019). Peptides are complex polymer chains combining (at least) twenty different amino acid monomers with different physico-chemical properties and thus, the combinatorial space is tremendous and scales by peptide length, *n*, as (at least) 20^n^. Although the specific mechanisms and prerequisites for potent peptide emulsifiers still remains only superficially characterized, our understanding of the underlying molecular properties continue to expand (Ricardo, Pradilla, Cruz, & Alvarez, 2021). Recent work has investigated the influence of factors such as interfacial peptide structure (Dexter, 2010; Du et al., 2020; García-Moreno et al., 2021; Lacou, Léonil, & Gagnaire, 2016), physico-chemical properties such as length and charge (García-Moreno, Gregersen, et al., 2020; Lacou et al., 2016; Liang et al., 2020; Yesiltas et al., 2021), amino acid composition (Enser, Bloomberg, Brock, & Clark, 1990; Saito, Ogasawara, Chikuni, & Shimizu, 1995; Siebert, 2001), and specific sequence patterns (Jafarpour, Gregersen, et al., 2020; Mondal et al., 2017; Nakai et al., 2004; Wychowaniec et al., 2020). Although various factors do appear to influence emulsification, the potential appears to indeed depend significantly on the propensity of a given peptide to adopt a more well-defined amphiphilic structure at the interface (Dexter & Middelberg, 2008; Enser et al., 1990; Saito et al., 1995). This property, in turn, is governed by these underlying factors. Although not appearing to be governed by the exact same molecular mechanisms, the stabilization of the air-water interface in foams have been suggested to also depend on peptide amphiphilicity (Enser et al., 1990; Jafarpour, Gregersen, et al., 2020).

Consequently, identification and molecular characterization of isolated peptides could enhance the understanding of functional mechanism and potential of enzymatic protein hydrolysates in food systems, such as emulsions and foams. Moreover, it would allow for development of targeted processes for release of specific peptides with known functional properties. This, in turn, could result in improved modification of a potato by-product to generate more value added ingredients with promising beneficial properties (Karami & Akbari-adergani, 2019). Waglay and Karboune, (2016a) characterized the structure of enzymatically generated peptides from potato protein, however, these authors did not correlate the specified characterization with functionality properties (Waglay & Karboune, 2016a). García-Moreno et al. (2020) investigated the emulsifying activity of six potato peptides (23–29 amino acids) predicted by bioinformatics as having potentially different predominant structure at the oil/water interface (e.g. α-helix, β-strand or unordered). The authors found that γ-peptides (half-hydrophobic and half-hydrophilic peptides with axial amphiphilicity), showed higher emulsifying activity, compared to α-helix and β-strand peptides, in agreement with their predictions (García-Moreno, Jacobsen, et al., 2020). However, generalization is not possible on such limited data. The study was followed by a more elaborate investigation of potato protein derived emulsifier peptides (García-Moreno, Gregersen, et al., 2020), showing that this could indeed not be generalized. In fact, the most promising peptide emulsifier (γ1) was later shown to adopt a predominantly α-helical conformation at the interface, thereby possessing both axial and facial amphiphilicity (García-Moreno et al., 2021). Although the structure-function relationship of emulsifier peptides from potato protein is more complex than predictable secondary structure and amphiphilicity, the two factors can be regarded as good indicators of emulsification potential (García-Moreno, Gregersen, et al., 2020).

Until now, most studies on protein hydrolysates are conducted using a trial-and error approach, where various industrial proteases are used to digest proteins in an untargeted manner. In such studies, process parameters (e.g. protease selection, pH, temperature, protein concentration, enzyme/substrate ratio, and time) are usually optimised in respect to bulk hydrolysate characteristics such as yield or functionality, with little or no attention to peptide-level insight. The application of mass spectrometry has in these instances mainly been focused on identification of peptides with high intensities in the bulk hydrolysate or in the high activity fractions. While such analysis may provide insight on peptides potentially responsible for the observed bulk functionality, it does not provide sufficient evidence unless functional properties are validated for the isolated peptide. In this study, we present the fundamentally different approach of data-driven, targeted hydrolysis. Building on existing knowledge on potato protein-derived peptide emulsifiers, we present a workflow where *in silico* sequence analysis is used as a guide for protease selection. By prediction of peptide release based on protease specificity, we hypothesise that application of specific proteases should produce a hydrolysate with better surface active (i.e. emulsifying and potentially foaming) properties. The approach is benchmarked against a range of commonly used industrial proteases, and the hydrolysates are characterised for their bulk physico-chemical and functional properties with particular focus on emulsification of fish oil and foam formation. Ultimately, we apply mass spectrometry-based proteomics analysis to qualitatively and quantitatively characterize the peptidome of the hydrolysates and relate these findings to both *in vitro* functionalities, predicted peptide release, and *a priori* knowledge on potato peptide emulsifiers. With this approach, we showcase how proteomics and bioinformatics may lay the basis for targeted process design in the future of peptide-based functional food ingredient development and production.

## 2. Materials and methods

Potato protein isolate (PPI) (87% protein, determined by Kjeldal-N and Dumas) was supplied by KMC AmbA (Brande, Denmark). The PPI was obtained using a proprietary, cold extraction method yielding native, non-denatured proteins. Alcalase 2.4L (2.4 AU/g), Neutrase 0.8L (0.8 AU/g), Flavorzyme 1000 L (1000 LAPU/g), and Trypsin (Pancreatic Trypsin Novo (PTN) 6.0S (6.0 AU/g) were provided by Novozymes A/S, (Bagsværd, Denmark). Distilled deionized water was used for the preparation of all solutions during hydrolysate production. As reference for emulsification experiments, sodium caseinate (SC) and purified, native patatin was used. SC (Miprodan 30) was supplied by Arla Foods Ingredients AmbA (Viby J, Denmark) and patatin was purified from the PPI by Lihme Protein Solutions (Kongens Lyngby, Denmark) using a gentle, sequential precipitation through a proprietary pH-shift methodology. All chemicals used were of analytical grade.

### 2.1. Target peptide selection, in silico sequence analysis, and process design for targeted hydrolysis

Previously investigated peptides derived from potato proteins were evaluated for emulsification potential based on published data (García-Moreno, Gregersen, et al., 2020; García-Moreno, Jacobsen, et al., 2020; García-Moreno et al., 2021; Yesiltas et al., 2021). Peptides were categorized (Table A.1) on a three-level scale (high (1), intermediate (2), and low (3)) according to their ability to i) reduce oil/water interfacial tension (IFT), ii) decrease oil droplet size, and iii) lead to physically stable emulsions during storage in comparison to SC. For IFT, peptides were evaluated based on their IFT at 30 min and classified as high if IFT < 15 mN/m, intermediated if 15 mN/m < IFT < 20 mN/m, and low if IFT > 20 mN/m (IFT for SC was reported as 10-14 mN/m in the four studies). For oil droplet size, peptides were evaluated by their mean diameter after emulsion (5% fish oil in water stabilized by 0.2% (W/V) peptide) production, either by D_4,3_ or D_3,2_ depending on what was reported in the individual studies. Peptides were classified as high if D < D_SC_, intermediate if D_SC_ < D < 3x D_SC_, and low if D > 3x D_SC_. For physical stability, peptides were classified as high if D did not increase by more than 100% after storage (compared to itself and/or SC) and no or little creaming was observed, intermediate if D did not increase by more than 300% after storage (compared to itself and/or SC) and middle/high creaming was observed, and low if D increased more than 300% after storage (compared to itself and/or SC) and/or severe creaming or separation was observed.

Peptide classified as high or intermediate in all three categories were selected for further *in silico* sequence analysis and ranked by their mean score across the three categories (and different studies, where applicable). To investigate potential release by enzymatic hydrolysis using the available proteases, the specific peptide was localized in the protein of origin (according to the original study) and the region of the protein containing the peptide and 15 amino acids up- and downstream (Table 1) extracted from Uniprot (Consortium et al., 2021). Cleavage specificity of the proteases was used to manually analyze potential hydrolysis of the protein region, where Tryp specificity is well-established and cleaves after Lys/Arg (K/R). Alc and Neut are broad specificity proteases but supplier specificity (Novozymes A/S) was used for *in* silico analysis. As such, Alc has a strong preference to cleave after Leu/Phe/Tyr/Gln (L/F/Y/Q), while Neut shows preference to cleave before Leu/Ile/Phe/Val (L/I/F/V). Flav is a complex mixture of endo- and exoproteases (Rabe et al., 2015), and cleavage specificity has not been established. As such, Flav was used merely as a reference protease due to its widespread use in the food industry. Based on distribution of target amino acids in and around the peptides, application of the individual proteases was evaluated, and the best possible process determined. To validate the approach and benchmark the method of targeted hydrolysis, all proteases were applied experimentally.

**Table 1:**
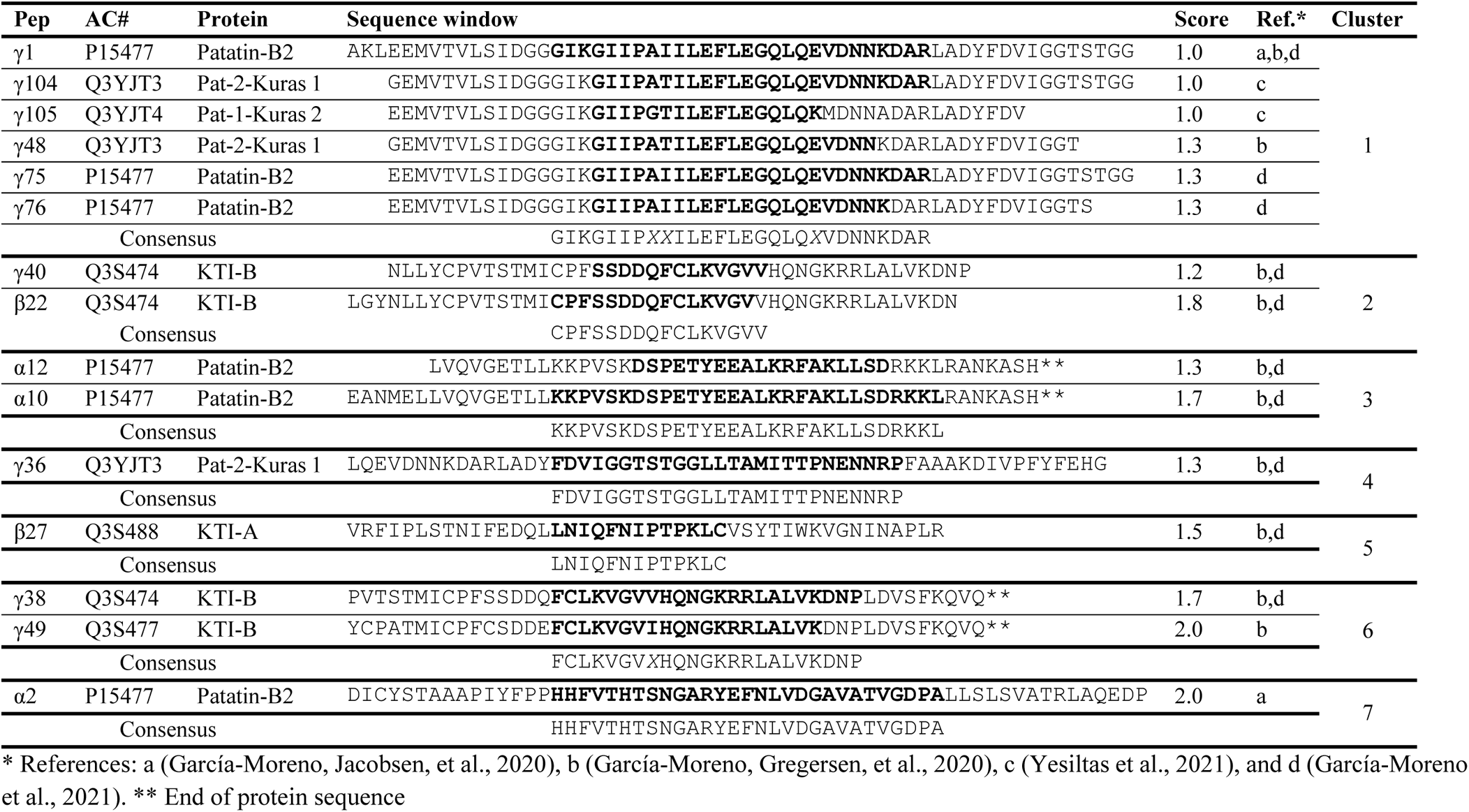
Cluster representation of final target peptides listing the peptides annotation from the reference study (Ref) along with the Uniprot identifier (AC#), protein name, sequence window (target peptide **(in bold)** along with the N- and C-terminal 15 amino acids cleavage window for *in silico* sequence and release analysis), average score (see Table A.1), and the cluster number (based on sequence overlap in identical or isoform proteins). Below each cluster, the aligned cluster consensus sequence is indicated with variable residues depicted as *“X” in italics*.

### 2.2. Potato protein hydrolysate preparation

Potato protein hydrolysates (PPHs) were produced from a native PPI, using two enzymatic hydrolysis strategies; (A) free-fall pH hydrolysis of native PPI, and (B) free-fall pH mode with protein heat denaturation prior to hydrolysis. In method A, a 1% (w/v) protein solution was prepared by gradual addition of PPI to distilled water and solubilized for 15 min by magnetic stirring. In method B, a 10% (w/v) PPI solution was prepared, heated to 90°C for 30 min, and resulting slurry was diluted 1:1 with distilled water to a final protein concentration of 4.35% (w/v). The pH of all PPI solutions was adjusted to 8.0 by 1M NaOH followed by addition of protease (Alcalase (Alc), Neutrase (Neut), or Flavourzyme (Flav), and Trypsin (Tryp)) at varying enzyme/substrate (E/S) ratio. For method A, E/S ratios of 0.1%, 0.5%, and 1% were applied while for method B, E/S ratios of 0.1% 1%, and 3% were applied. In both methods, hydrolysis was carried out at 50°C for 2 h. pH and temperature was selected to accommodate activity ranges of all proteases according to manufacturer (Novozymes A/S supplied information).

Following hydrolysis, the pH of the solution was adjusted to 7.0 with either 1M NaOH or 1M HCl and supernatants were heated to 90 °C for 15 min for enzyme inactivation. After cooling by tap water, solutions were centrifuged at 10,000 ×g for 20 min at 20 °C and the supernatant collected. The PPH was lyophilized and stored at 4 °C until further analysis. The two applied hydrolysis strategies are illustrated in Fig. 1.

**Fig. 1:**
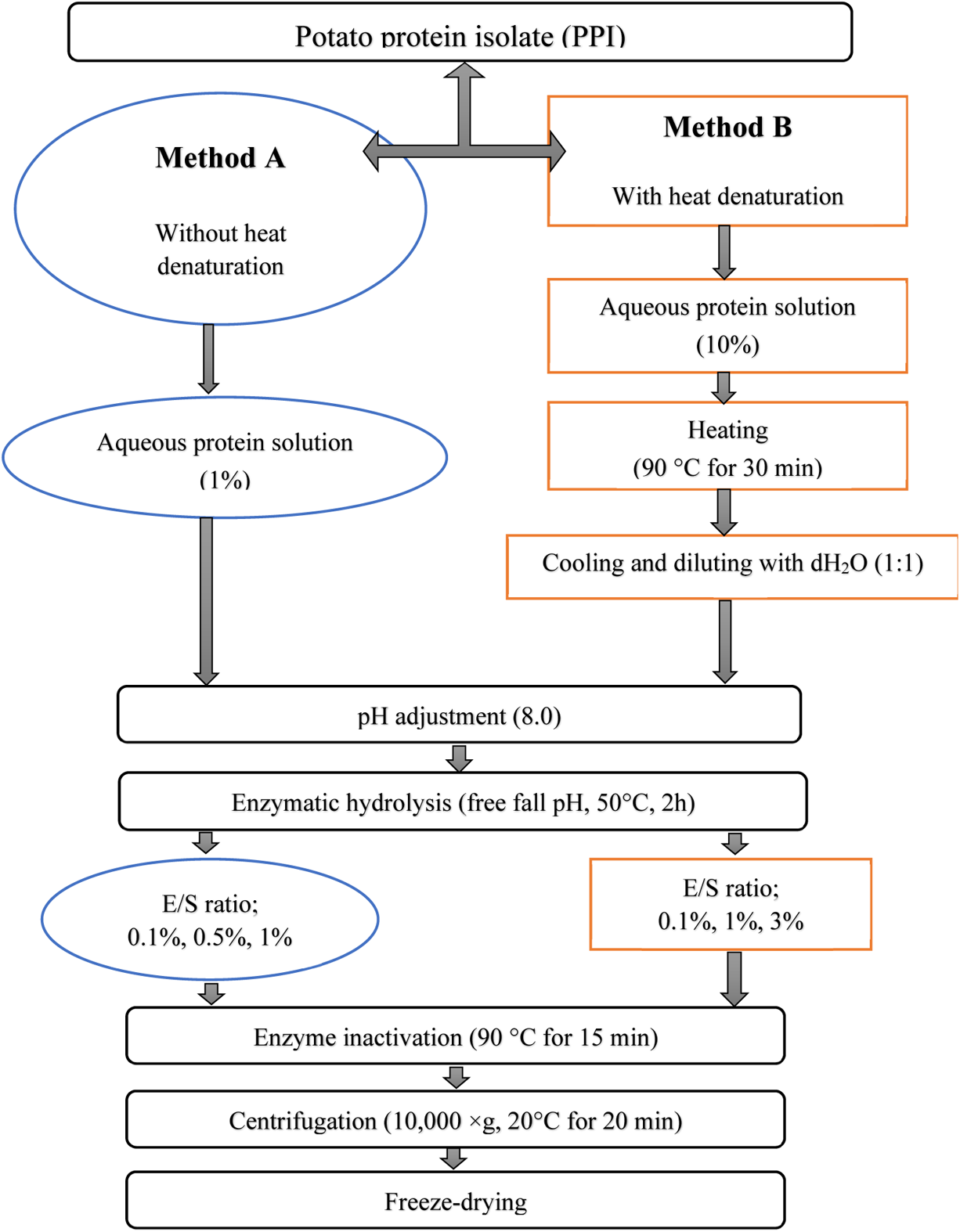
Flow chart for the enzymatic hydrolysis of potato protein isolate (PPI) by application of Alc, Neut, Flav or Tryp using two different processes; (A, left) without heat denaturation of PPI and (B, right) with heat denaturation to inactivate protease inhibitors. Steps specific for method A are depicted in circles while steps specific for method B are depicted in squares. Common steps are depicted as rounded squares spanning both workflows.

### 2.3. Degree of hydrolysis and peptide chain length

Degree of hydrolysis (DH) was determined based on α-amino nitrogen content as previously described (Jafarpour, Gregersen, et al., 2020) with minor modifications, adjusting for the α-amino nitrogen content of the untreated substrate. Briefly, free α-amino content of PPHs was determined using the PFAN-25 free amino nitrogen assay kit (PractiChrom, USA) measured using A_530_ on a Picoexplorer (USHIO INC, USA), according to manufacturer guidelines, using glycine as reference for standard curve generation. DH was calculated as:

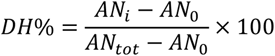

where *AN_i_* is the concentration of α-amino nitrogen (mM/g substrate) resulting from hydrolysis at a given time, *i*, *AN_0_* is the α-amino nitrogen content of the untreated substrate, and *AN_tot_* is the total amount of α-amino nitrogen content following complete hydrolysis with 6 M HCl at 110°C for 24 h, as previously described (Jafarpour, Gomes, et al., 2020). *AN_tot_* was based on duplicate amino acid (AA) analysis and calculated using the molecular weight of individual AAs. All determinations of α-amino-N were performed in triplicates.

The determined DH for the individual PPHs was used to estimate the average peptide chain length (PCL_DH_) as:

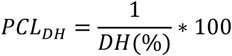

### 2.4. Nitrogen recovery and protein content

Total nitrogen content was determined using the Dumas combustion method using a fully automated Rapid MAX N (Elementar Analysensysteme GmbH, Langenselbold, Germany), and nitrogen recovery (NR) in the soluble fraction was determined as previously reported (Jafarpour, Gomes, et al., 2020) and calculated as:

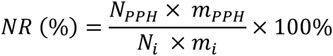

where *N_PPH_* is nitrogen content (%) in the PPH, *m_PPH_* is the mass (g) of analysed PPH, *N_i_* is the nitrogen content (%) of the initial substrate, and *m_i_* is the mass (g) of the initial substrate analyzed. Prior to analysis, the system was calibrated using multiple blanks, aspartic acid, and wheat protein isolates. The protein content was estimated using a standard industrial nitrogen to protein conversion factor of 6.25 (Jones, 1931; Mariotti, Tomé, & Mirand, 2008)

### 2.5. Yield

The yield of hydrolysis was determined as the mass ratio to the initial substrate mass, as

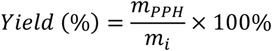

where *m_PPH_* is the mass (g) of obtained PPH after lyophilization and *m_i_* is the mass (g) of initial substrate.

### 2.6. PPH solubility

The relative solubility at 10 mg/mL of PPI/PPHs was determined based on nitrogen content by Dumas, as previously described (Jafarpour, Gomes, et al., 2020). Briefly, 200 mg PPH was dissolved in 20 mL of 0.1 M sodium phosphate buffer (pH=7.4), vortexed for 10 s, shaken at 80 rpm for 30 min, and centrifuged at 7500 ×g for 15 min. PPH solubility was calculated as:

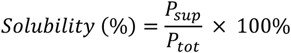

where *P_sup_* is protein/peptide content in supernatant and *P_tot_* is total protein/peptide content in PPI the respective PPH as obtained for determination of NR. Measurements were performed in triplicate.

### 2.7. Bulk Density

The bulk density of the PPHs was determined according to (Jafarpour, Gomes, et al., 2020). Briefly, 5g PPH was added to a 50 mL graduated cylinders and gently tapped 10 times on the lab bench. Bulk density was reported as g/mL.

### 2.8. Color parameters

The tristimulus color parameters (*L*a*b\**) of PPHs were recorded using a Miniscan XE colorimeter (Hunter Lab, Reston, Virginia, USA) and the whiteness of PPHs was determined in accordance with (Hashemi & Jafarpour, 2016) and calculated as:

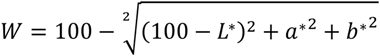

Where, *W* is whiteness index, *L*\*; indicating lightness from black (0) to white (100); *a*\*; indicating redness from green (−120) to red (+120); and *b*\*; indicating yellowness going from blue (−120) to yellow (+120). Measurements were performed in triplicate.

### 2.9. Emulsifying properties

Emulsifying activity index (EAI) and emulsion stability index (ESI) were determined using the method described by (Pearce & Kinsella, 1978) with slight modifications, previously described (Jafarpour, Gregersen, et al., 2020). Briefly, 15 mL PPH in distilled water (2 mg/mL) was mixed with 5 mL rapeseed oil by an ultraturax homogenizer (IKA, Germany) at speed of 9,500 rpm for 60 s without pH adjustment. Fifty μL aliquots were pipetted from the bottom of the container at 0 and 10 min after homogenization and added to 5 mL of 0.1% sodium dodecyl sulfate (SDS) solution and mixed by gentle shaking. The absorbance of the diluted solution was measured at 500 nm using a spectrophotometer (SHIMADSU UV-1280, Japan). EAI was calculated as:

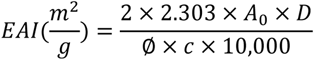

where *A_0_* is the absorbance at 500nm immediately following homogenization, *D* is dilution factor (100), *Ø* is oil volume fraction (0.25) and *c* is protein concentration (g/mL).

ESI was calculated as:

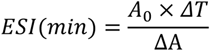

where ΔT is equal to 10 min and ΔA is the difference in absorbance at 500 nm after 0 and 10 min (ΔA=A_0_-A_10_). Enriched patatin, untreated PPI, and SC solutions (2mg/mL) were used as references. Measurements were performed in triplicates.

### 2.10. Foaming properties

Foaming capacity (FC) and foaming stability (FS) were determined according to the method of (Elavarasan, Naveen Kumar, & Shamasundar, 2014) and calculated as previously described (Jafarpour, Gregersen, et al., 2020). Accordingly, 0.1 % (w/v) PPH in distilled water was homogenized at 9,500 rpm for 120 s using an Ultraturrax (IKA, Germany), and poured into a 200 mL graduated cylinder. FC was calculated as:

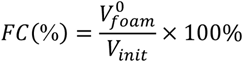

where *V^0^_foam_* is the foam volume immediately after homogenization and *V_init_* is the initial sample volume.

FS was determined after a 30 min resting period (FS_30_) and calculated as:

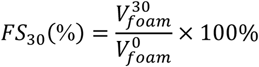

where *V^0^_foam_* is the foam volume after 30 min. Measurements were performed in triplicates.

### 2.11. 1D SDS-PAGE analysis

For visualization of the hydrolysis using both strategies, PPHs were analyzed by SDS-PAGE. using SurePAGE 4-20% gradient Bis-Tris gels (Genscript, Picastaway, NJ, USA) under reducing conditions, as previously described (Jafarpour, Gregersen, et al., 2020). As controls, the unhydrolyzed PPI sample and a process control (PPI with no added protease but same treatment) were included.

### 2.12. Peptide analysis by LC-MS/MS

Lyophilized PPH was solubilized in a detergent-containing buffer, reduced and alkylated in-solution, and desalted using C-18 StageTips, and dried by SpeedVac, as previously described (Jafarpour, Gomes, et al., 2020). Desalted peptides were solubilized in solvent A (0.1% aq. formic acid (FA)) and 1µg (determined by NanoDrop (Thermo Scientific Bremen, Germany)) was loaded and separated on an EASY-nLC (Thermo Scientific) equipped with a reverse phase (RP) Acclaim Pepmap Nanotrap column (C18, 100 Å, 100 μm.×2 cm, nanoViper fittings (Thermo Scientific)) followed by a RP Acclaim Pepmap RSLC analytical column (C18, 100 Å, 75 μm.×50 cm, nanoViper fittings (Thermo Scientific). Eluted peptides were introduced into a Q Exactive HF mass spectrometer (Thermo Scientific) via a Nanospray Flex ion source (Thermo Scientific) using a fused silica needle emitter (New Objective, Woburn, MA, USA). Samples were loaded at 8 μL/min and eluted by constant flow at 300 nL/min during a 120 min ramped gradient, ranging from 5 to 100% of solvent B (0.1% formic acid (FA), 80% (V/V) acetonitrile). MS data was acquired in positive mode using a Top-20 data-dependent method. Survey scans were acquired from 200 m/z to 3,000 m/z at a resolution of 60,000 at 200 m/z and the HCD fragmentation spectra were acquired at a resolution of 30,000 at 200 m/z using an isolation window of 1.2 m/z and a dynamic exclusion window of 30 s. The maximum ion injection time was set to 150 ms for both MS and MS/MS scans. ACG target was set to 3e6 and 2e5 for MS and MS/MS scans, respectively. Charge exclusion was only applied for unassigned isotope peaks (all charge states allowed). Peptide match and exclude isotopes were enabled.

### 2.13. LC-MS/MS data analysis in MaxQuant

MS raw data was analyzed in MaxQuant v.1.6.10.43 (Cox & Mann, 2008; Tyanova, Temu, & Cox, 2016), as previously described (Jafarpour, Gomes, et al., 2020). Briefly, unspecific *in silico* digestion was employed to identify peptides in the range 3-65 AAs. Data was searched against a manually curated version of the full protein database for *Solanum tuberosum* (tax:4113) from UniProt (Consortium et al., 2021) where redundant fragments were removed, as previously described (García-Moreno, Gregersen, et al., 2020). Standard settings were applied, using a 5% false discovery rate (FDR) on both peptide and protein level (Gregersen et al., 2022), including reverse sequences for FDR control, and including common contaminants.

### 2.14. Summary statistics and mean peptide properties

Following removal of false positives and contaminants, Venn diagrams were created to visualize peptide identifications and similarity between PPHs. Peptide tables were treated according to the intensity-weighted peptide abundance estimation methodology, as previously described (Jafarpour, Gomes, et al., 2020; Jafarpour, Gregersen, et al., 2020), and average peptide length (PCL_avg_) was calculated as:

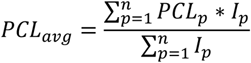

where *PCL_p_* is the length of peptide *p* of *n* identified and quantified peptides and *I_p_* is the MS1 intensity of the same peptide.

### 2.15. Qualitative and quantitative correlation with known potato peptide emulsifiers

For correlation of identified peptides with known emulsifier peptides derived from potato proteins, a set of seven selected target clusters were constructed (Table 1). Here, single AA substitutions were disregarded. For each cluster, a representative cluster sequence spanning the entire sequence with “X” representing variable residues (i.e. AAs where substitutions occur within the cluster peptides) was created. All peptides related to the class/family of proteins associated with the cluster were initially included and subjected to filtering based on two criteria. To be regarded as a potentially emulsifying peptide contributing to the bulk activity observed, the identified peptides were >12 AAs in length, which was previously identified as the minimum length to have emulsifying properties (García-Moreno, Gregersen, et al., 2020; Yesiltas et al., 2021). In addition, a sequence overlap of any peptide of >50% with the representative cluster sequence was required. To facilitate this, all lead proteins (i.e. the first protein in the protein group constructed in MaxQuant) within a relevant class/family of proteins (i.e. patatins and all protease inhibitors) were aligned using Clustal Omega (Sievers et al., 2011). Aligned proteins within each class/family were grouped into protein sequence clusters where a specific target cluster sequence was represented. Within each protein sequence cluster, sequence identity was not required but AA numbers were aligned. Sequence overlap between identified peptides and a representative cluster sequence was determined through the following set of equations:

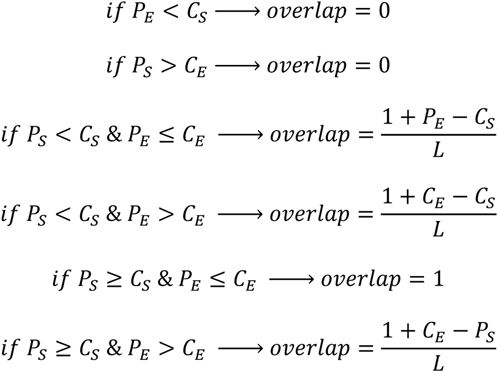

where P_S_ and P_E_ are the AA positions for the start and end of an identified peptide when mapped to a certain protein cluster sequence, C_S_ and C_E_ are the AA positions for the start and end of a representative cluster sequence when mapped to the same protein cluster sequence, L is the length of the identified peptide. Peptides were classified as >50%, >75%, and >95% overlap, thereby with increasing confidence of emulsifier activity with increasing sequence overlap. The five most abundant (by MS1 intensity) peptides from each PPH with >50% overlap with representative cluster sequence were extracted and mapped to the representative cluster sequence using the NCBI blastp suite. The alignment was then visualized in the NCBI multiple sequence alignment (MSA) viewer using the built-in hydropathy color scale and with substitutions indicated.

### 2.16. Protein-level quantitative distribution of PPH peptides

To determine the protein-level distribution of peptides in PPHs, the method of unspecific, length-normalized relative intensity, I_L_^rel^, was used (Gregersen et al., 2021, 2022). In short, the relative molar abundance of an identified protein (group) was estimated as:

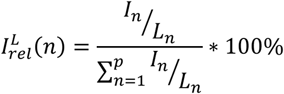

where I_n_ is the intensity of protein n (i.e. sum of all peptide intensities ascribed to the protein (group)) of p quantified proteins in a given PPH and L_n_ is the length of protein n. Subsequently, proteins were grouped according to family/class (García-Moreno, Gregersen, et al., 2020), and the relative abundance of each class was determined.

### 2.17. Statistical Analysis

The statistical significance between measurements was determined by variance analysis (ANOVA) using Statgraphics software (version 18.1.06 for Windows), and means were compared by Duncan’s multiple comparison post-test. Statistical differences were considered to be significant at p <0.05.

## 3. Results and Discussion

### 3.1. In silico analysis and design of a targeted hydrolysis process

Based on previously published data on *in vitro* emulsifying properties of potato protein-derived peptides, 58 peptides were evaluated and ranked by their ability to reduce oil/water interfacial tension, mean droplet size after emulsification of 5% fish oil in water, and physical stability of the emulsion (Table A.1). From the initial evaluation, 15 unique peptides with strong emulsifying properties were selected for further *in silico* analysis to identify release potential by enzymatic hydrolysis (Table 1). The 15 selected peptides group into seven clusters by sequence similarity, where particularly cluster 1 is densely populated. Here it is evident that cluster peptides are all related to γ1, being either truncated variants or truncated isoforms of the peptide.

Alc has broad specificity and has been reported to cleave after a wide range of residues (Ala/Leu/Val/Phe/Tyr/Trp/Glu/Met/Ser/Lys) with varying claims throughout literature (Doucet, Otter, Gauthier, & Foegeding, 2003; Lei, Cui, Zhao, Sun-Waterhouse, & Zhao, 2014; Lu et al., 2021; Sbroggio, Montilha, Figueiredo, Georgetti, & Kurozawa, 2016). A similar lack of consensus for Neut specificity is also found, although varying reports on the broad specificity of Alc and Neut exist, an *in silico* analysis based on supplier specificity (Novozymes A/S) was performed. Using the target peptide sequence and including a 15 amino acid N- and C-terminal cleavage window, all cleavage sites of Alc, Neut, and Tryp were mapped (Fig. 2).

**Fig. 2:**
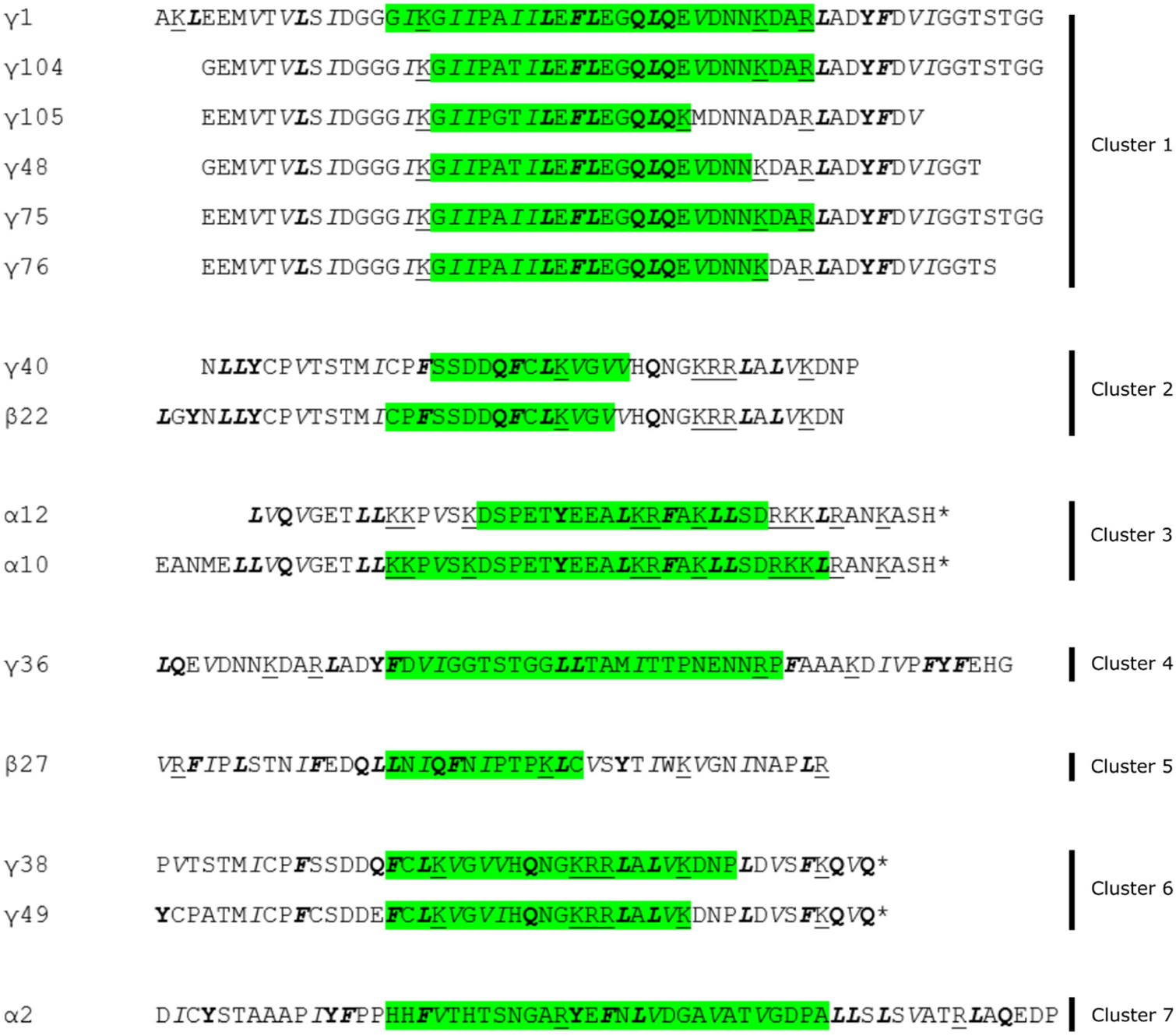
*In silico* analysis of release potential for selected target peptides following enzymatic hydrolysis by Tryp, Alc, or Neut. Peptides with validated *in vitro* emulsifying properties (highlighted in green) are listed with a 15 amino acid N- and C-terminal cleavage window and clustered by sequence similarity (Table 1). Cleavage sites for Tryp (cleavage after R/K) are <u>underlined</u>, cleavage sites for Alc (cleavage after L/F/Y/Q) are highlighted in **bold**, and cleavage sites for Neut (cleavage before I/L/F/V) are highlighted in *italics*. Alc and Neut both cleave at Phe (F) and Leu (L). “*” indicates the end (C-terminus) of the native protein sequence.

In cluster 1, a good overall compatibility with tryptic cleavage is observed. Tryp hydrolysis will result in a three AA N-terminal truncation of γ1 (resulting in γ75) and a three AA C-terminal truncation of γ75 (resulting in γ76). As the C-terminal Lys in γ76 is followed by an Asp, cleavage efficiency for Tryp may be reduced in this position (Giansanti, Tsiatsiani, Low, & Heck, 2016), making γ75 the most probable product, as previously reported (García-Moreno, Gregersen, et al., 2020; García-Moreno et al., 2021). While γ48, γ105, and γ106 originate from other isoforms of patatin and thus contains different AA substitutions, placement of tryptic AAs are highly favourable as well. Target AAs for both Alc and Neut are abundant within all cluster 1 peptides making release of the peptides less likely. Nevertheless, target AAs for Neut are located favourably in both termini of cluster 1 peptides, and some release of highly related peptides may be possible if excessive internal hydrolysis is avoided, although this is considered unexpected in a controlled and reproducible manner.

In cluster 2, target AAs for Tryp are unfavourably distributed. Similarly to cluster 1, target AA for both Alc and Neut are abundant in both peptides, although partial hydrolysis could release peptides very similar to the targets for both proteases. In cluster 3, α12 is fully embedded in α10. Nevertheless, both peptides have centrally located target AAs of all three proteases, making them unlikely products of hydrolysis. However, placement of AAs in the terminal regions may make them decent targets for partial hydrolysis by all three proteases. Cluster 4 contains a single peptide (γ36) and is found in the region of patatin immediately following cluster 1. Hydrolysis with Tryp would introduce a four AA N-terminal elongation and a potential one AA truncation in the C-terminal, although the Pro may also reduce efficiency (Giansanti et al., 2016). Although target AAs for both Alc and Neut are found through the peptide, the positioning of Phe residues at the termini makes γ36 a good target for partial hydrolysis. This may particularly be the case for Alc, as there are substantially fewer target AAs for this protease in the sequence. In clusters 5-7, target peptides have unfavourable positioning of target AAs for all three proteases, but partial hydrolysis by Alc and Neut may be possible for obtaining peptides closely resembling the target.

Ultimately, the *in silico* analysis shows that particularly cluster 1 is an excellent target for hydrolysis with Tryp. Tryp may also produce a peptide closely resembling γ36 from cluster 4. For the remaining clusters, no clear evidence for release of target (or closely related peptides) was found, although both Alc and Neut may produce hydrolysates containing peptides resembling the targets through partial hydrolysis, where Alc may produce peptides with better emulsifying properties than Neut.

### 3.2. Enzymatic hydrolysis

Initially, enzymatic hydrolysis was performed without heat denaturation due to the high (90%) protein solubility (Table 2) in the native PPI (method A). However, a very low efficiency of hydrolysis was observed. The low degree of hydrolysis was observed in the lyophilized supernatant following post-hydrolysis heat inactivation of the proteases and subsequent centrifugation (Fig. 3A), by comparison to the control (unhydrolyzed PPI). Only for high concentrations of Flav (Fig. 3A, lane 12) and to a lesser extent Alc (Fig. 3A, lane 9), some degree of hydrolysis of the patatin band (∼40 kDa) could be observed. This can be ascribed to the maintained inhibitory activity of wide variety of protease inhibitors inherent to potato tubers (Kunitz-type ∼20 kDa, PIN-type ∼15 kDa, and MCPI-type ∼5-10 kDa) It has been suggested that in a temperature range of 55-70 °C, inhibitor activity of protease inhibitors in potato protein is remarkably destroyed (Van Koningsveld et al., 2001). However, it has also been shown that even after cooking of potato protein at high temperature (75-100 °C), around 10% of the chymotrypsin inhibiting activity remains (Huang, Swanson, & Ryan, 1981). Application of harsh treatments, such as combination of thermal coagulation and acid precipitation, may therefore destroy inhibitory activity of some protease inhibitors (including aspartate-, cysteine-, and Kunitz-type protease inhibitors), while other protease inhibitors may retain their inhibitory function (Waglay & Karboune, 2016a). This is particularly of interest, as PTN 6.0S has been reported to exhibit chymotrypsin activity (Nongonierma, Paolella, Mudgil, Maqsood, & FitzGerald, 2017). Wang and Xiong (2005) investigated hydrolysis of heat-denatured potato protein by SDS-PAGE, and after 30 min Alc hydrolysis, the ∼40 kDa band vanished as a result of patatin hydrolysis (Wang & Xiong, 2005). Based on this, we hypothesized that the efficiency of enzymatic hydrolysis will be enhanced if the protease inhibitors become thermally denatured and hence, method B was employed. Although increased hydrolysis is desired, the degree of hydrolysis (DH) should be kept low, as peptides should be above a certain length to retain emulsifying properties (García-Moreno et al., 2016). The minimum length depends on peptide interfacial conformation (García-Moreno, Gregersen, et al., 2020), but at least 12 AAs. This directly implies that, the DH should not exceed 10% and preferably be even lower (Klompong, Benjakul, Kantachote, & Shahidi, 2007; Liu, Kong, Xiong, & Xia, 2010; Tamm et al., 2015). Consequently, the hydrolysis time was kept short (2 h) for method B.

**Fig. 3:**
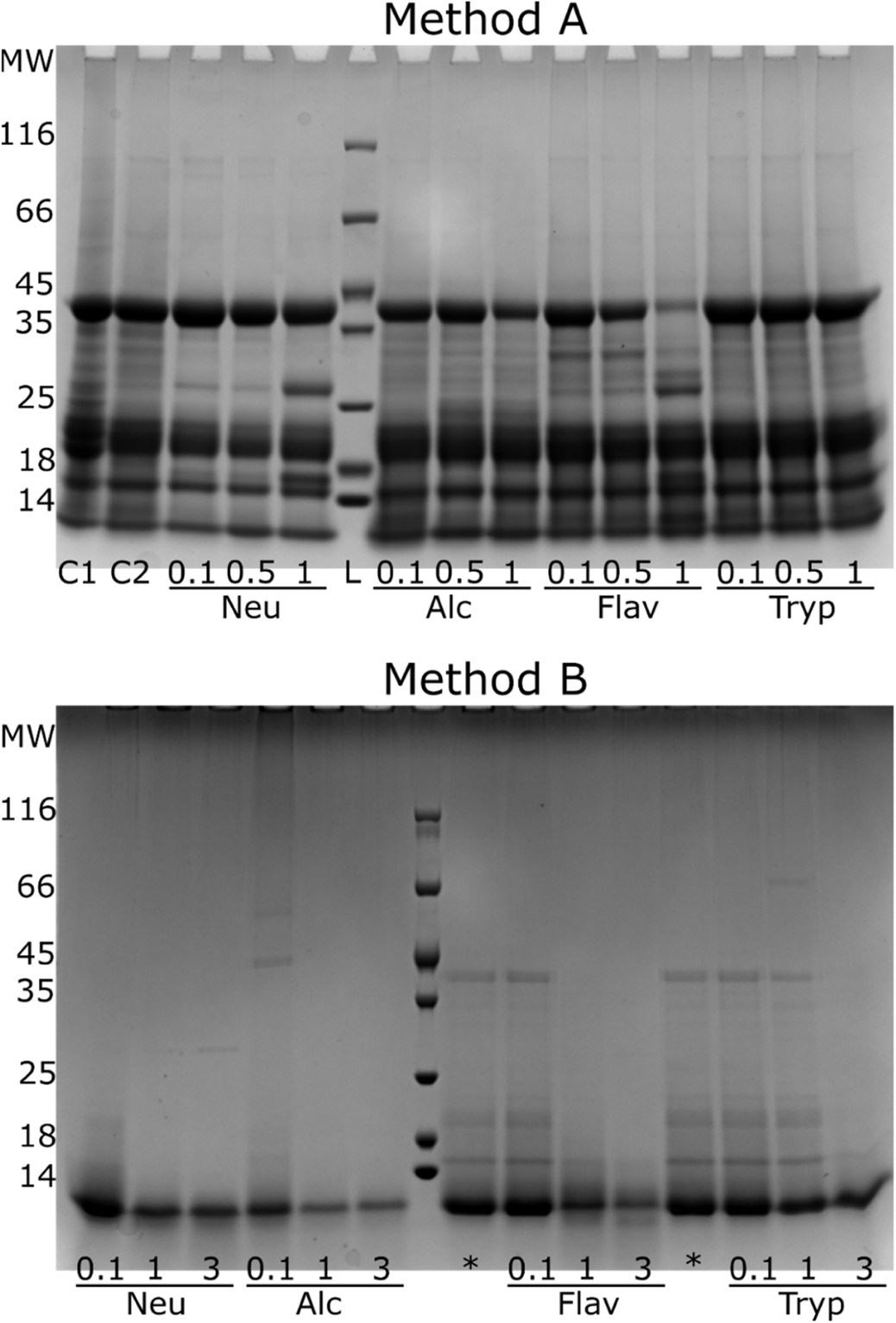
SDS-PAGE of PPH (freeze-dried supernatants) following hydrolysis of native PPI (Method A) and hydrolysis of heat denatured PPI (Method B) using Neutrase (Neu), Alcalase (Alc), Flavourzyme (Flav), or Trypsin (Tryp) at different E/S ratios (Method A: 0.1%, 0.5%, and 1%. Method B: 0.1%, 1%, and 3%). C1: Native PPI. C2: Process control with native PPI. *Repetition of 0.1% hydrolysis for Flav and Tryp.

**Table 2:**
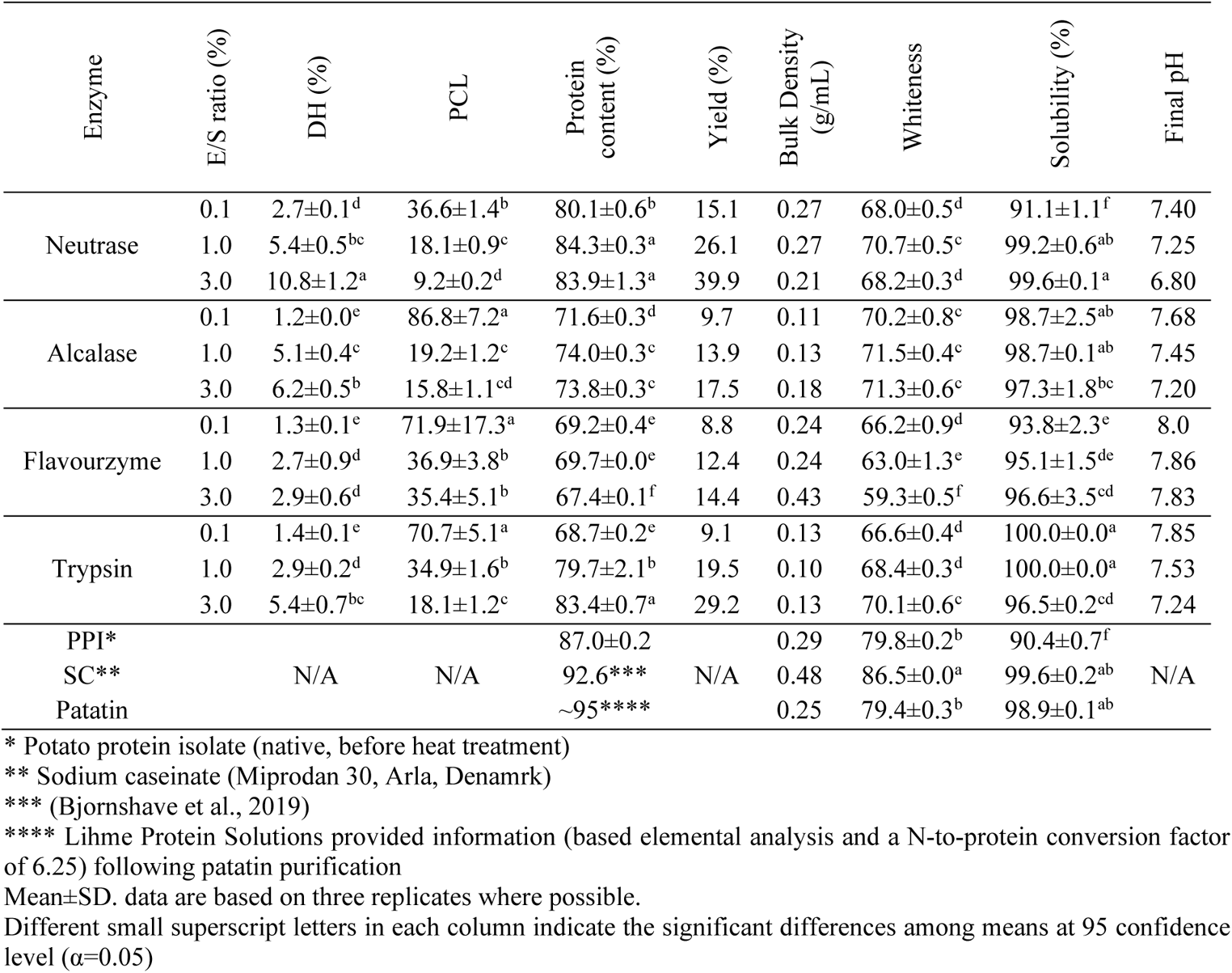
Hydrolysis and bulk properties of enzymatic potato protein hydrolysates (PPH)* by different industrial proteases at different E/S ratios. For each PPH, the degree of hydrolysis (DH) by α-amino-N determination and the associated average peptide chain length (PCL) are indicated along with protein content (% by N*6.25), hydrolysis yield (%mass in freeze-dried supernatant), bulk density, whiteness, and solubility (at 10 mg/mL). For comparison, bulk properties for the initial substrate (potato protein isolate, PPI), sodium caseinate (SC), and a purified patatin fraction are listed.

Enzymatic hydrolysis was conducted in a free fall pH mode, i.e., the initial pH was adjusted at 8.0, but during hydrolysis gradually decreased at varying extent to a final pH of 6.8-8.0 for method B (Table 2). This was done to emulate industrial scale-up, where pH control may not always be possible (Kamnerdpetch et al., 2007). As expected, a relation between the extent of hydrolysis (DH) and the decrease in pH was observed. All hydrolysates displayed DH within the desired range (DH<8%) with the exception of 3% Neut showing significantly (p<0.05) higher extent of hydrolysis (DH=11%). Lower DH for Flav has also been observed in other studies investigating proteolysis of plant proteins (Li et al., 2015; Zhao et al., 2012). As expected, increasing E/S ratio increased the efficiency of enzymatic reaction significantly (p<0.05), but at different extents. This is supported by SDS-PAGE analysis (Fig. 3B), where a dramatic decrease of unhydrolyzed protein, compared to the non-denatured PPI (Fig. 3A), can be observed particularly for higher E/S ratios. This is in agreement with previous studies, where with fixed, initial substrate concentration and fixed hydrolysis time, increased E/S ratio increases DH (Waglay & Karboune, 2016a).

Higher proteolytic activity for Flav, compared to e.g. Alc (DH of 22% vs. 8%), has previously been reported for hydrolysis of potato pulp and attributed to the simultaneous endo- and exoproteolytic activity of Flav (Kamnerdpetch et al., 2007). Nevertheless, as the study employed substantially higher E/S ratio (7%), prolonged hydrolysis (26 h), and was performed directly on pulp, direct comparison is futile. Wang and Xiong (2005) investigated the hydrolysis of heat-denatured potato protein using Alc (1% E/S ratio) and stated that by increasing the reaction time from 0.5 h towards 1h and 6h, DH increased from 0.72 to 1.9 and 2.3%, respectively. These results are comparable to the DH obtained using Alc in our study. Although Alc is often reported to result in higher DH due to broader specificity (Demirhan, Apar, & Özbek, 2011; García Arteaga, Apéstegui Guardia, Muranyi, Eisner, & Schweiggert-Weisz, 2020; Jafarpour, Gregersen, et al., 2020; O’Keeffe & FitzGerald, 2014), a significantly higher (p<0.05) DH is observed for Neut. This despite the initial pH of the hydrolysis is better aligned with optimum conditions for Alc. The lower DH for Alc is, however, in line with previous studies on potato protein hydrolysis (Kamnerdpetch et al., 2007; Wang & Xiong, 2005). Because a substantial amount of protease inhibitory activity is retained even after heating (Van Koningsveld et al., 2001), the mode of action for the applied proteases (i.e. protease families/classes) and the composition of protease inhibitors in PPI becomes a limiting step. Alc is a serine protease in the subtilisin family (Aldred, Phang, Conlan, Clare, & Vancso, 2008; Donlon, 2007) while Neut is a Zn-metalloprotease (Wu & Chen, 2011) but both exhibit endoproteolytic activity. As previously reported (García-Moreno, Gregersen, et al., 2020), protease inhibitors constitute more than half of the protein in the PPI (referred to as KMC-Food) used as substrate for hydrolysis. The vast majority of these are Kunitz-type inhibitors, where the class of serine protease inhibitors (KTI-B) constitute a very large proportion, corresponding to around 20% of the total protein (García-Moreno, Gregersen, et al., 2020; Pęksa & Miedzianka, 2021). In contrast, metalloprotease inhibitors constitute a very small part (>0.1%) and, importantly, are all in the form of metallocarboxypeptidase inhibitors (MCPI), thereby inhibiting exoproteolytic activity (García-Moreno, Gregersen, et al., 2020; Jørgensen et al., 2011; Pouvreau et al., 2001). Consequently, retained inhibitory activity against e.g. serine protease like Alc may likely explain why a significantly higher DH is observed for Neut. Interestingly, DH also appears to be somewhat correlated with both protein content in the PPH and the yield of hydrolysis (Table 2), indicating that hydrolysis is indeed a prerequisite to resolubilise the heat-denatured PPI, in line with previous studies (Miedzianka et al., 2014; Wang & Xiong, 2005). The significantly lower DH (p<0.05) observed for Flav is also reflected in significantly lower (p<0.05) protein content and lower yields at all E/S ratios. Low yields (<20%) and a substantial reduction in protein content (<80%) compared to PPI (87%) is also observed for Alc and Tryp at low E/S ratios (0.1 and 1 %), thereby making such processes potentially unbeneficial from an industrial and economical point of view.

### 3.3. Bulk properties of PPH

#### 3.3.1. Physical properties

The bulk density has a substantial impact on e.g. packaging and handling attributes, as well as physicochemical and sensory properties of powdery-like materials such as freeze-dried protein hydrolysates (Kurozawa, Morassi, Vanzo, Park, & Hubinger, 2009; Rodríguez-Díaz, Tonon, & Hubinger, 2014). In our study, PPH produced using Neut and Flav display bulk densities (0.21-0.27 g/mL) within the same range as the native PPI and the enriched patatin fraction (Table 2). These values are a little lower than previous reports (0.36 g/mL) for freeze-dried potato protein (Claussen, Strømmen, Egelandsdal, & Strætkvern, 2007) but comparable to e.g. casein and codfish hydrolysates (Jafarpour, Gomes, et al., 2020; Sarabandi, Sadeghi Mahoonak, Hamishekar, Ghorbani, & Jafari, 2018).

Color parameters of recovered PPHs were measured using the CIE system and reported as lightness (*L*\*), redness (*a*\*), and yellowness (*b*\*) (Table A.2) as well as the overall vector of these three parameter colors, whiteness (Table 2). PPHs showed lower whiteness values compared to the PPI and patatin powders (P<0.05). Among PPHs, the whitest (70.2-71.5) powder was obtained by application of Alc, whereas Flav resulted in the lowest whiteness (59.2); particularly at highest E/S ratio (P<0.05). Miedzianka et al. (2014) reported that application of Alc enzyme for hydrolyzation of fodder potato protein significantly lightened the color of PPH powder (72.5) compared to the raw sample (50.3) (Miedzianka et al., 2014). In this study, the PPI had a substantially higher whiteness (79.8) whereas the whiteness of PPHs were comparable to that reported by Miedzianka et al. (2014). If decreased whiteness in PPHs is related to the heat treatment prior to hydrolysis, and thereby possibly a result of enzymatic browning prior to enzyme denaturation (Sapers & Miller, 1992), oxidation (Tien, Vachon, Mateescu, & Lacroix, 2001), or Maillard reactions (Keppler, Schwarz, & van der Goot, 2020; Liska, Cook, Wang, & Szpylka, 2016), was not explored further. However, the enzyme formulation may itself have direct impact on the PPH color (Fig. A.1), as the Flav formulation was substantially darker than the remaining formulations (Fig. A.2) and the whiteness decreased with increasing E/S ratio. Similar observations have been reported for hydrolysates from round scad (Thiansilakul, Benjakul, & Shahidi, 2007).

#### 3.3.2. Emulsifying Properties

Turbidity measurements to quantify EAI and ESI are considered reliable indicators of emulsifying properties for proteins and peptides (Pearce & Kinsella, 1978). The EAI and ESI of PPHs prepared with the four proteases are shown in Table 3. In general, application of lower E/S ratios (0.1% and 1%) resulted in significantly lower EAI compared to controls (PPI, SC and Pat) (P<0.05), and only Tryp 1% displayed EAI comparable to PPI and SC (P>0.05) but still significantly lower than native Pat (P<0.05). In contrast, the EAI of Tryp PPHs at 3% E/S ratio was significantly higher (P<0.05) compared to both PPI and SC and comparable to the EAI of native Pat (P>0.05), confirming the original hypothesis based on *in silico* analysis. 3% Alc PPH also displayed EAI higher than PPI and SC (P<0.05) but was significantly lower than both Pat and 3% Tryp PPH (P<0.05). At 3% E/S ratio, Flav treatment resulted in a nearly equivalent (P>0.05) EAI compared to those from PPI and SC, while Neut PPH displayed the lowest EAIs (P<0.05). That isolated, native Pat has strong emulsifying properties is in line with previous studies (Van Koningsveld et al., 2006). Using other methods to evaluate emulsifying properties than EAI (e.g. emulsion droplet size distribution and oil/water interfacial tension reduction) has, however, revealed that native Pat may not result in such strong emulsification as observed here, whereas the 3% Tryp PPH does appear to indeed have very strong emulsifying properties regardless of evaluation method (Manuscript submitted, Food Chemistry). In any case, obtaining native, isolated patatin is a costly process, and therefore not a scalable and economically viable solution.

**Table 3:**
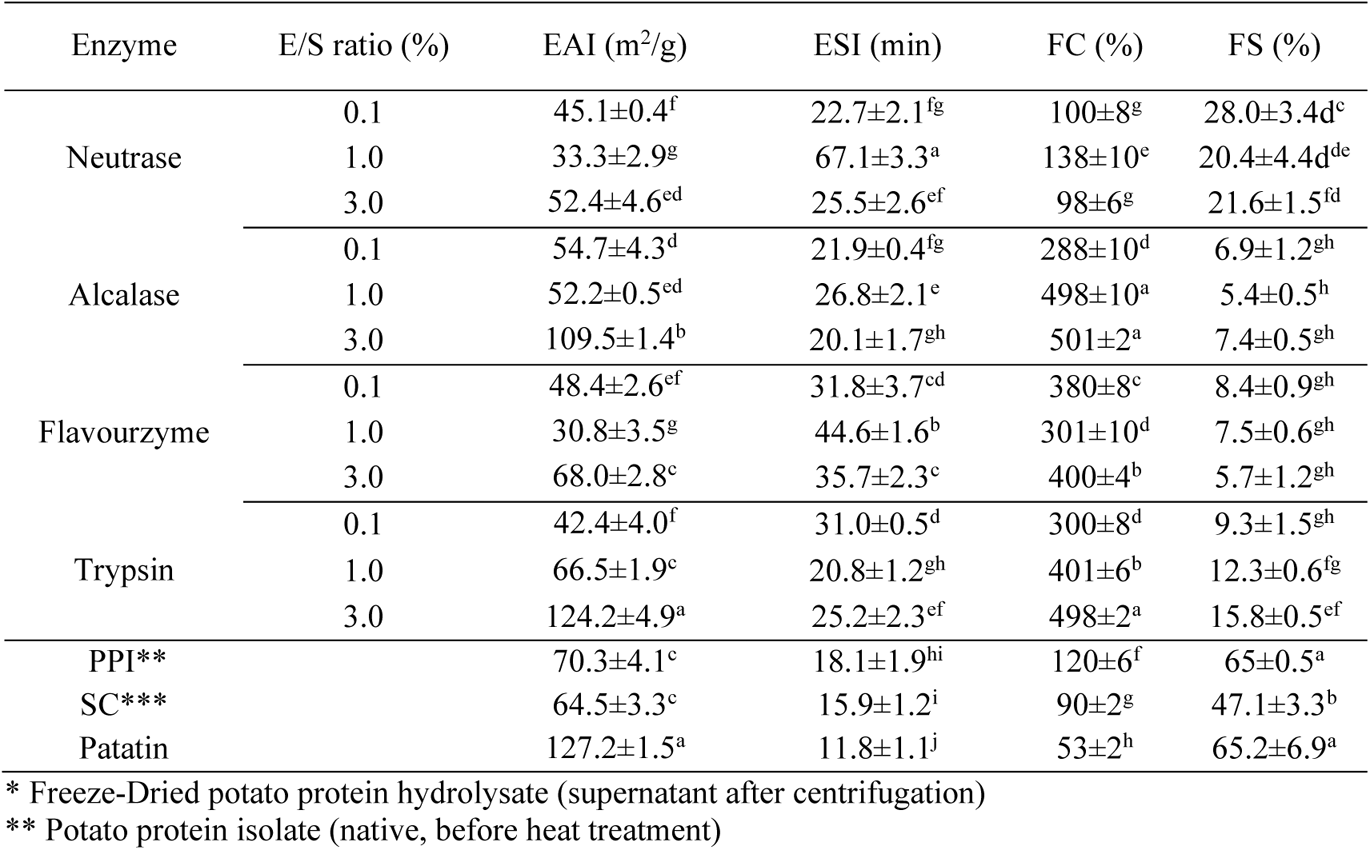
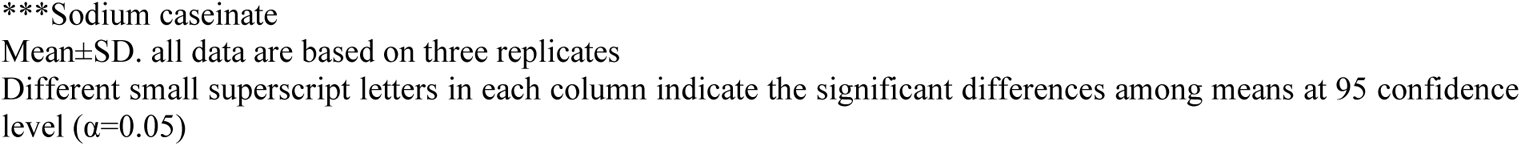
Emulsifying and foaming properties of enzymatic potato protein hydrolysates (PPH)* by different industrial proteases at different E/S ratios. For each PPH, the Emulsification Activity Index (EAI), Emulsification Stability Index (ESI), Foaming Capacity (FC), and Foaming Stability (FS) are indicated. For comparison, the native potato protein isolate (PPI), sodium caseinate (SC), and a purified patatin fraction are listed.

All PPHs in our study produced emulsions with higher ESI compared to those with PPI, SC and Pat (P<0.05). The highest ESI value was determined for Neut (67.0 min) followed by Flav (44.5 min), both at 1% E/S. However, these PPHs also displayed the lowest values of EAI across all PPHs. EAI does not follow the same trend as a function of E/S ratio (and thus DH) for the investigated proteases. For instance, in case of Neut and Flav by increasing the E/S ratio from 0.1% to 1%, the EAI values decreased, whereas it increased significantly at 3% E/S (P<0.05). On the other hand, the EAI of Tryp derived PPH, showed a constantly increasing trend with increasing enzyme concentration, reaching 124 m^2^/g at 3% E/S (corresponding to DH = 5.4%). EAI values of Alc derived PPH showed no significant difference between 0.1 and 1% E/S ratio (P>0.05), whereas it increased significantly (P<0.05) to 109 m^2^/g at 3% E/S (DH = 6.2%). In line with these results, Zhao and Hou (2009) reported that soy protein hydrolysates produced with Tryp (DH 1– 2%) exhibited a better EAI than those of hydrolyzed by Neut at the same DH (X. Zhao & Hou, 2009). This could be ascribed to the higher specificity of Tryp, resulting in a more well-defined hydrolysate and release of peptides containing a hydrophilic C-terminus (Lys/Arg), which could contribute to electrostatic stabilization of the oil droplets in the emulsion (X. Zhao & Hou, 2009). Similarly, other studies revealed the superior solubility and emulsifying properties of protein hydrolysates produced with Tryp (Padial-Domínguez, Espejo-Carpio, Pérez-Gálvez, Guadix, & Guadix, 2020; Taherian, Britten, Sabik, & Fustier, 2011).

Overall, EAI and ESI did not correlate with DH nor each other directly (Fig. A.3). Diffusion properties in solution highly influence the adsorption rate of emulsifiers to the oil-water interfaces during homogenization (McClements & Jafari, 2018). In other words, there is a propensity for smaller monomers to diffuse more quickly to an interface than a e.g. larger proteins or aggregates. Despite a significantly higher DH for Neut at 3%, the EAI was less than half of the EAI for Alc and Tryp derived PPHs (corresponding to DH = 6.2% and DH = 5.4%, respectively). In contrast, Flav-derived PPHs displayed much lower DH (2.9%) but higher EAI than Neut PPHs. This indicates that although smaller peptides diffuse rapidly and adsorb at the interface, they may indeed be less efficient in stabilizing emulsions. Moreover, the results also indicate that mean peptide chain length (PCL) as determined by the DH cannot be used directly as a measure of emulsification, as previously shown in potato protein hydrolysates (Akbari, Mohammadzadeh Milani, & Biparva, 2020). Previous studies have indicated that a minimum length (>12 AAs) is required for a peptide to obtain a defined structure at the interface and thus display emulsifying properties (García-Moreno, Gregersen, et al., 2020), indicating that there may be a preferred length range for peptides to display emulsifying properties. Although there are no clear trends in our data, it does appear that operating within a range of fairly low DH (∼ 2-8%) seems to promote the likelihood of obtaining a hydrolysate with improved emulsification, when evaluated by EAI and ESI (Fig. A.3.A and Fig. A.3.B), but this also appear to be highly protease-dependent and not universally applicable. Interestingly, there are also indications, but no strong evidence, that there may be a trade-off between EAI and ESI for the hydrolysates (Fig. A.3.C). Although DH (and thus PCL) is not a viable measures to estimate emulsifying activity by itself, it may be one of many factors to consider; particularly in relation to avoiding excessive hydrolysis, which can substantially deteriorate emulsifying activity (Klompong et al., 2007; Liu et al., 2010; Vioque, Sánchez-vioque, Clemente, Pedroche, & Millán, 2000). The amphiphilicity and interfacial conformation of peptides may, ultimately, outweigh peptide length as a determining factor for emulsifying properties (Klompong et al., 2007).

Importantly, our results indicate that Tryp is indeed capable of producing a PPH with significantly improved emulsifying properties compared to both other industrially relevant proteases and the substrate itself. In fact, only the application of Tryp was able to improve both emulsification activity and stability significantly (P<0.05), compared to untreated PPI. These observations are in line with the predicted and expected outcome based on prior knowledge and *in silico* analysis, which indicates the potential release of known emulsifier peptides from potato protein when using Tryp. This highlights that a data-driven, targeted approach for enzymatic hydrolysis is a promising and viable approach for optimizing the functional parameters of industrial side streams such as potato protein. And that this may be obtained in a predictable manner, alleviating the need for conventional trial-and-error methodology.

#### 3.3.3. Foaming properties

A common way to evaluate the foaming properties of products such as hydrolysates, is by determinination of their foaming capacity (FC) and stability (FS), describing different molecular properties related to stabilization of the air/water interface (Petruccelli & Anon, 1995; Ralet & Guéguen, 2000). In our study, apart from Neut-derived PPHs at 0.1% and 3% E/S ratio, all PPHs showed significantly higher (P<0.05) FC compared to control samples. In contrast to the emulsifying properties, the highest FC of the control samples was determined for PPI, followed by SC, and patatin, respectively (P<0.05). The PPHs from Alc and Tryp showed remarkably high (∼500%) but comparable (P>0.05) FC at high E/S ratio (3%) (Table 3). These results imply that there is no direct relationship between foaming capacity and DH (Fig. A.4.A). Conformational properties of released peptides relies highly on the specificity of the applied protease. For instance, the lower FC as well as EAI of Neut-derived PPH may be attributed to a sequence-specific and lower surface activity of released peptides rather than their average length; similarly as was observed for emulsifying properties. That being said, the decrease in FC with a decrease in E/S ratio from 1% to 3% may also be the result of extensive hydrolysis, ultimately releasing too short peptides with a decreased propensity to form defined structures at the interface. This is in agreement with previous studies on Alc hydrolysis of a potato protein isolate, where extending hydrolysis time, increased DH up to 17%, which resulted in a decrease in FC (Akbari et al., 2020). Such high DH was not observed for Alc (or other) PPHs in this study, which could explain why the effect was only seen for Neut-derived PPHs as all other PPH had a DH in the range from 1.2% - 6.2% (Table 2).

Control samples showed higher (P<0.05) stability compared to PPHs (Table 3), as the highest FS (∼65%) was determined for the native PPI and the patatin fraction (P>0.05), followed by SC at ∼47% (P<0.05). For PPHs, the highest FS was determined for Neut followed by Tryp PPHs, while both Alc and Flav PPHs presented the lowest FS (<10%) among all (P<0.05). Increasing the Neut E/S ratio from 0.1% to 3% caused a significant decrease (P<0.05) in stability of formed foams from 28% to about 22% (P<0.05), whereas the inverse effect was observed for Tryp, where increasing E/S from 0.1% to 3% significantly increased FS from ∼9% to ∼16% (P<0.05). Between Alc and Flav PPHs, there was no significant difference (P>0.05) in determined FS. Interestingly, only hydrolysis by Tryp improved both FC and FS (as well as EAI) with increasing DH. In earlier studies, the FC and FS of a potato protein concentrate (8% and 5.3%, respectively) increased to 162% and 51% after hydrolysis by Alc for 2h (Miedzianka et al., 2014). Although these values are not in full agreement with our data, the study also highlights the positive effect of proteolysis for increasing the surface activity, as well as solubility, compared to the protein substrate. In our study, Alc and Tryp hydrolysis at 3% E/S ratio resulted in both comparable FC (∼500%) and DH (∼6%), but remarkably different FS. Foams produced with 3% Tryp PPH had double the stability of 3% Alc-derived PPH (Table 3). No apparent relation between DH and FS was observed (Fig. A.4.B)

Similarly to the relation between EAI and ESI, there appears to be a general trade-off between FC and FS (Fig. A.4.C), although e.g. Tryp PPHs shows the opposite trend. While control samples (PPI, SC, and Pat) produce the most stable foams, they produce the least stable emulsions. Hydrolysis of PPI (as well as native Pat) significantly increases the capacity to foam (P<0.05), but also significantly decreases the foam stability (P<0.05), in line with the general observations of a negative relation between FC and FS for PPHs in this study. Although no clear correlation is observed, there does appear to be some relation between high emulsifying and foaming capacities (Fig. A.5.A), indicating that to some extent, similar molecular properties are involved in both interfacial properties, in line with previous studies (Wouters, Rombouts, Fierens, Brijs, & Delcour, 2016). The stability of the interfaces, however, appear to be governed by different forces and properties, and no apparent relation (Fig. A.5.B).

### 3.4. Peptide identification and mean peptide properties

Across the 12 PPHs investigated with LC-MS/MS, unspecific analysis in MaxQuant resulted in identification of 46,316 unique peptides following removal of reverse and contaminant peptides. Although a higher FDR (5%) was applied for MaxQuant analysis, this level was previously shown to be suitable for non-specific digests due to the significantly increased combinatorial search space (Gregersen et al., 2022). In general, a lower number of peptides were identified in PPHs produced using Neut, although more than 10,000 peptides were identified in all PPHs (Table 4). This is a tremendous increase in depth of analysis compared to previous reports of LC-MS/MS analysis on hydrolysates, where the number of identified peptides is often reported in the tens to low thousands range (Caron et al., 2016; Cui, Sun, Cheng, & Guo, 2022; Hinnenkamp & Ismail, 2021; Y. P. Huang, Dias, Leite Nobrega de Moura Bell, & Barile, 2022; Jafarpour, Gregersen, et al., 2020; Jin, Yan, Yu, & Qi, 2015; M. Li, Zheng, Lin, Zhu, & Zhang, 2020).

**Table 4:** Summary statistics for identified peptides (Peptide IDs) and peptide weighted average length (PCL_avg_) by unspecific analysis of potato protein hydrolysate (PPH) LC-MS/MS data in MaxQuant. The total peptide MS1 intensity (Σ Int) for each PPH is listed along with the associated normalization factor (NF) used for relative, peptide-level comparison.

| Enzyme | Alcalase |  |  | Flavourzyme |  |  | Neutrase |  |  | Trypsin |  |  |
| --- | --- | --- | --- | --- | --- | --- | --- | --- | --- | --- | --- | --- |
| E/S ratio | 0.1% | 1% | 3% | 0.1% | 1% | 3% | 0.1% | 1% | 3% | 0.1% | 1% | 3% |
| Peptide IDs | 13140 | 14106 | 13125 | 14622 | 12936 | 13094 | 12210 | 11944 | 10829 | 14948 | 12756 | 12475 |
| Total | 19513 |  |  | 21976 |  |  | 17356 |  |  | 23509 |  |  |
| $PCL_{avg}$ | 15.9 | 15.3 | 14.9 | 15.8 | 14.8 | 14.5 | 14.2 | 13.2 | 12.7 | 15.9 | 15.8 | 15.0 |
| $\Sigma Int [1E^{12}]$ | 8.7 | 12.2 | 13.2 | 6.9 | 5.5 | 8.1 | 13.6 | 12.0 | 9.3 | 5.7 | 10.6 | 13.7 |
| NF | 0.638 | 0.889 | 0.966 | 0.502 | 0.399 | 0.589 | 0.996 | 0.877 | 0.683 | 0.420 | 0.773 | 1 |

As expected, changing hydrolysis conditions by increasing E/S ratio, and thereby increasing DH, had a substantial effect on the number and nature of identified peptides for all four investigated proteases. This is illustrated both by the number of identified peptides (Table 4) as well as shared peptides between PPHs obtained using the same protease (Fig. A.6). Interestingly, the effect of protease and E/S ratio on DH, determined through α-amino nitrogen quantification (Table 2), is not to the same extent reflected in the peptide level data, using intensity-weighted peptide abundance estimation, PCL_avg_ (Table 4). This is in contrast to previous studies, where a much stronger correlation between the two methods was observed (Jafarpour, Gregersen, et al., 2020). However, there are notable differences between the two studies. In Jafarpour et al. (2020), hydrolysis was performed on a raw side stream from the cod industry, where this study deals with a much purer protein isolate. This is clearly illustrated by the lack of distinct protein bands in SDS-PAGE analysis in the cod hydrolysates, where we here observed strong bands from intact protein and larger protein fragments by SDS-PAGE following hydrolysis (Fig. 3), indicating that a substantial amount of protein remains in forms undetectable using a bottom-up proteomics approach. According to Linderstrøm-Lang theory, proteases may have higher affinity for intermediate fragments/peptides than intact proteins (Adler-Nissen, 1986; Linderstrøm-Lang, 1953). It was previously shown in e.g. milk (Deng, van der Veer, Sforza, Gruppen, & Wierenga, 2018; Hinnenkamp & Ismail, 2021) and rice (Nisov, Ercili-Cura, & Nordlund, 2020) as well as potato (Akbari et al., 2020; Pȩksa & Miedzianka, 2014) proteins, that this is indeed the case in a highly protein- and protease-specific manner. This also suggests that the observation of intense bands for residual intact protein by SDS-PAGE (Fig. 3), should likely not be interpreted as lack of hydrolytic activity, but rather increased protease affinity for intermediate fragments/peptides. Similarly, hydrolysis to the single AA and dipeptide level will contribute significantly to the total DH of a sample, while these remain undetected in the MS experimental design. Nevertheless, a decrease in PCL_avg_ is observed with increasing E/S ratio for each protease, and substantially lower PCL_avg_ values are obtained for Neut corresponding to the higher DH in these PPHs. This shows that even in spite of the challenges imposed by intact protein and other undetected species, MS data and PCL_avg_ can provide an indication for the progression of hydrolysis in addition to peptide- and protein-level insight.

### 3.5. Identification of known peptide emulsifiers from potato proteins

Based on emulsifying properties of the PPHs, a deeper peptide-level analysis of peptides associated with the seven sequence clusters associated with known and highly potent emulsifiers (Section 3.1) was performed. As the major constituents of the potato proteome (i.e. patatin and protease inhibitors) all represent a large number of protein isoforms, mapping peptides to the isoforms is a challenging task. This is particularly the case as single AA substitutions and minor truncations/elongations may not have a detrimental effect of the peptide functionality (Enser et al., 1990; García-Moreno, Gregersen, et al., 2020; García-Moreno, Jacobsen, et al., 2020; Ricardo et al., 2021). To accommodate this, we established a workflow, where identified peptides were mapped onto representative target cluster sequences (Fig. 2) by defining lead protein sequence clusters (Table 1). The workflow allows for determining the degree of overlap between identified peptides and the representative cluster sequence while allowing for substitutions, truncations, and elongations. By using the peptide MS1 intensity as an estimate of abundance, it is thus possible to determine how much of the peptide MS1 intensity for the whole PPH is constituted by peptides adhering to both the length requirement and a minimum sequence overlap with a given target cluster. As the total MS1 intensity varies substantially between PPHs (Table 4), unhydrolyzed proteins content varied between PPHs, and the same total amount was loaded on-column (1µg), total MS1 intensities were normalized relative to the PPH with the highest MS1 intensity (Tryp 3%). After applying the normalization factors (Table 4), the normalized, relative MS1 intensity constituted by peptides with >50%, >75%, and >95% overlap with each of the seven target clusters, was determined for each PPH using both a 12 AA (Fig. 4, left) and 15 AA (Fig. 4, right) minimum length requirement. From here, we see that peptides mapping to target peptide clusters 1 and 3 account for the largest contributions of mapped peptides regardless of required sequence overlap and length requirement. We also see that peptides mapping to clusters 2, 5, and 6 have practically no contribution to the sum, while the contribution from cluster 4 and 7 peptides is low. Moreover, we observe a shift in which PPHs has the highest relative contribution to overlapping peptides, based on the requirement of degree of overlap.

**Fig. 4:**
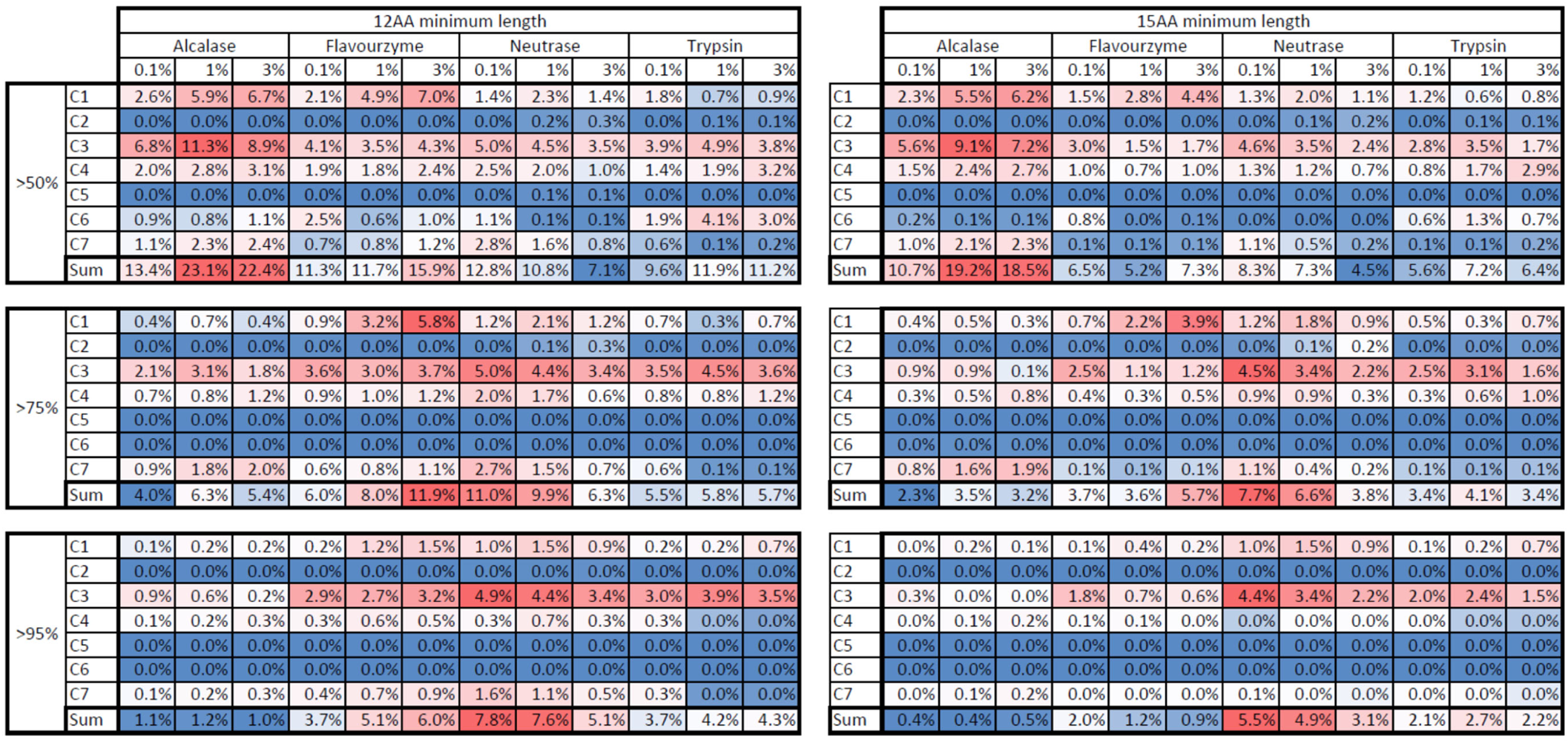
Heat maps (blue (low) to red (high)) of quantitative (by MS1 I_rel_) sequence overlap between identified peptides and the target cluster sequence for all seven target cluster (C1-C7). Overlaps are given with increasing minimum overlap (50%, 75%, 95% (top to bottom)) and increasing minimum peptide length (12 AAs (left) and 15 AAs (right)) requirements. Sums across all clusters for each condition are color coded separately from the individual clusters to illustrate overall adherence of PPH peptides to all target clusters.

Interestingly, target cluster overlap (Fig. 4) overall shows low agreement with observed emulsifying properties of the PPHs (Table 3). While a low degree of overlap (>50%) shows that Alc PPHs (particularly at 1% and 3% E/S which also show high EAI) has the highest proportion of overlapping peptides, increasing requirement of sequence overlap shift the highest proportion of overlapping peptides towards Flav and Neut PPHs (which show lowest EAI), while Alc PPHs here show the lowest proportion. In all cases, Tryp PPHs (showing highest EAI), show an intermediate proportion of overlapping peptides in comparison and a low content of cluster 1 peptides, which were the primary target by Tryp hydrolysis. This observation led us to investigate if certain regions of a target cluster may be more important. By extracting the sequences for high intensity peptides in each PPH with >50% overlap in cluster 1 and 3 (Table A.3), it is possible to see that Neut PPHs contain abundant peptides overlapping with cluster 1, but that all are located in the C-terminal region of the cluster (Fig. 5, left). In contrast, high intensity Alc peptides are found in the N-terminal region of the cluster. While high intensity Flav peptides are located in both cluster termini, Tryp PPH peptides, particularly at 3% E/S, span more of the cluster (Fig.5, left). This indicates that the N-terminal region of the cluster may be of higher importance, and, more importantly, that peptides also should cover at least a certain part of the cluster to attain the interfacial activity. This is in agreement with previous studies (Yesiltas et al., 2021), where cluster 1 peptides (e.g. γ105) are highly truncated in the C-terminal region of the target cluster sequence. As such the region covered by γ105 (GIIPGTILEFLEGQLQK) may be regarded as the core region of cluster 1 and could represent a “critical region” for emulsifying activity, as this region produces a highly amphiphilic α-helix at the interface. With the exception of γ1 (which is cleaved by trypsin after Lys in position 3 resulting in γ75), all cluster 1 peptides (full length and/or full length isoforms) were identified in the 3% Tryp PPH (Table A.4). Some were also identified in the 1% Tryp PPH, while very minute amounts (<0.002% I_rel_) were found in 3% Flav PPH, both verifying that target peptides were indeed released in a targeted manner and that observed differences in emulsifying activity may be ascribed to substantial presence of these highly functional peptides. Nevertheless, this type of analysis only describes a subpopulation of the entire, complex PPH, and does not account for all other peptides and their potential (positive or negative) contribution to the bulk functionality of the PPHs. Our results also indicate that a quite substantial amount of Tryp was needed to efficiently release the peptides, which may be ascribed to residual inhibitory activity in the PPH despite heat treatment.

**Fig 5:**
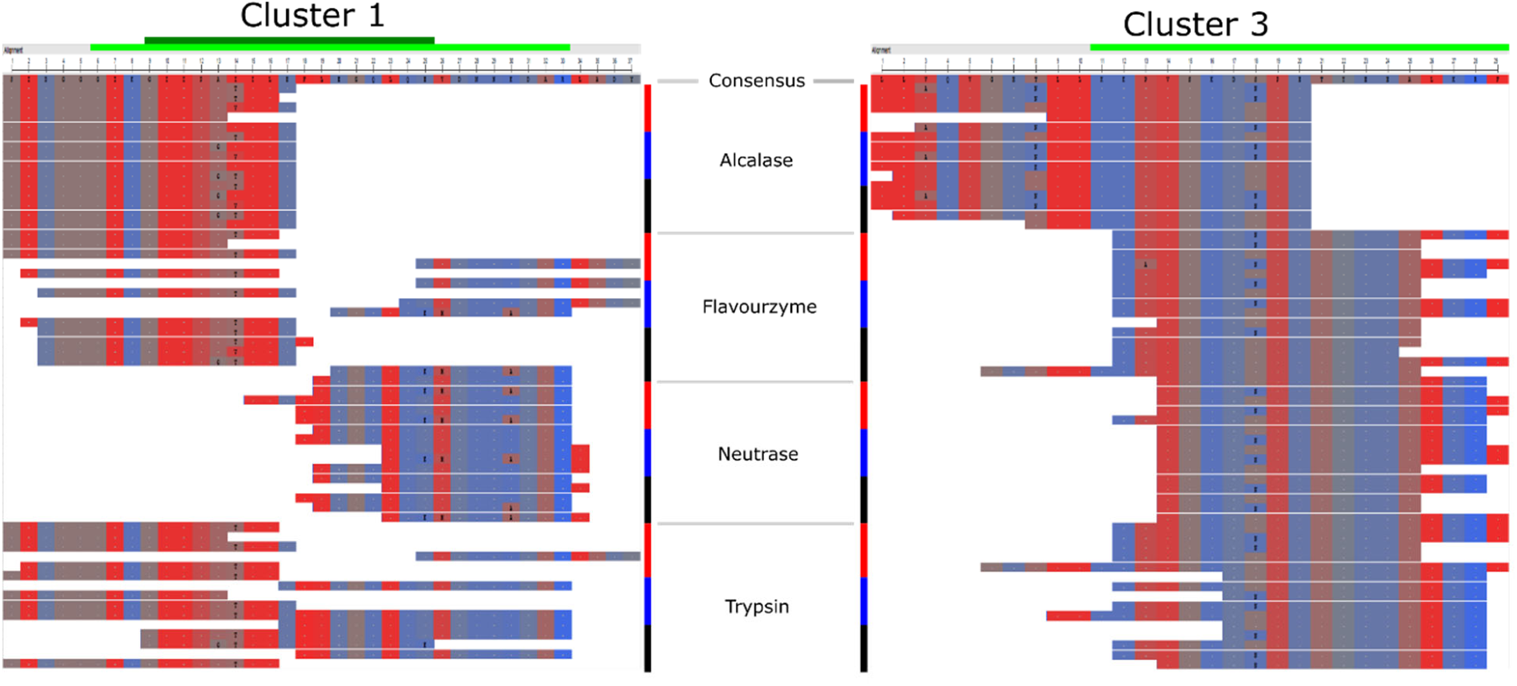
Alignment of top-5 high intensity peptides in Alc, Flav, Neut, and Tryp (top to bottom) PPHs for 0.1% (red), 1% (blue), and 3% (black) E/S ratio. Only peptides with at least 50% overlap with the target cluster consensus sequence for Cluster 1 (left) and Cluster 3 (right) are shown. For each condition (protease and E/S ratio), the top-5 peptides are depicted with descending relative MS1 intensity (top to bottom). Amino acids are color coded from red (hydrophobic) to blue (hydrophilic) according to the NCBI MSA Viewer hydropathy color scale. The consensus sequence for each cluster is shown by a light green bar (top), while the suggested “core region” for cluster 1 peptides is shown in dark green. The consensus sequence was extended in both termini to allow for full mapping and visualization of all top-5 overlapping peptides. Single amino acid substitutions (relative to the consensus sequence) are assigned by the substituent single letter code on the individual peptide level.

Cluster 3 peptides constitute the majority of all overlapping cluster peptides. With increasing requirements for both length and degree of overlap, the highest proportion shifts from Alc PPHs to Flav and particularly Neut and Tryp PPHs (Fig. 4). Cluster 3 represents two patatin-derived peptide emulsifiers, α10 and α12, which were previously shown to indeed adopt a helical conformation at the oil/water interface (García-Moreno et al., 2021). High intensity Flav, Neut, and Tryp peptides all appear to cover a significant amount of the target cluster sequence (Fig. 5, right), which intuitively should make all these PPHs good emulsifiers. Nevertheless, if assuming a helical conformation with 3.6 AA per turn, the distribution of AAs in these peptides do not appear favourable for producing an amphiphilic helix. Particularly high intensity Neut peptides are slightly truncated in the N-terminal region of the cluster, making the characteristic pattern of AA distribution in an amphiphilic helix (Eisenberg, Weiss, & Terwilliger, 1982; García-Moreno, Gregersen, et al., 2020; Yesiltas et al., 2021) absent, to a large degree. In contrast, Alc peptides extend N-terminally of the consensus sequence in cluster 3. This means that peptides include a region, which have a highly favourable AA distribution for adopting a highly amphiphilic helical conformation at the interface, and also forms an amphipathic helix in native patatin (Fig. A.7). Furthermore, the most abundant cluster 3 peptides in Alc PPHs (Fig. 5, right) are variants of the same peptide (LLAQVGENLLKKPVSKDNPE), containing either single AA substitutions or minor (1-2 AA) N-terminal truncations, which are likely to not have substantial effect on the interfacial properties. Furthermore, most of these peptides have a highly hydrophobic N-terminus, similarly to the cluster 1 core sequence, which may serve as a hydrophobic anchor to facilitate stronger adsorption to the oil/water interface and high emulsion capacity. As Tryp and, to a lesser degree, Flav peptides also extend into this region compared to Neut peptides, this may be a key part of why Alc PPHs also show good emulsifying properties and why Flav PPHs perform better than Neut PPHs.

Although the contribution from cluster 4 peptides is smaller than cluster 1 and 3 peptides, it is noteworthy that a peptide (LADYFDVIGGTSTGGLLTAMITTPNENNRPFAAAK), corresponding to a slightly elongated version of γ36 (FDVIGGTSTGGLLTAMITTPNENNRP), was identified in all Tryp PPHs, exactly as predicted (Section 3.1). With increasing E/S ratio, the estimated abundance of this peptide also increased from 0.08% (I_rel_) in 0.1% Tryp to 0.64% (I_rel_) in 3% Tryp (Table A.4). This in spite of the total MS1 intensity more than doubled in 3% Tryp, indicating that increasing the DH towards completion for tryptic hydrolysis, substantially increases the release of the target peptide. This also correlates well the high emulsifying and foaming properties observed for 3% Tryp PPH. The peptides was surprisingly also identified in all Alc and Flav PPHs, but at a substantially lower abundance (I_rel_ < 0.05%). In Alc PPHs, several peptides of sufficient length for helical surface activity (>15 AAs) and covering the most of γ36, were identified at noteworthy I_rel_ (Table A.4). This is particularly the case in 3% Alc, where five peptides (16-34 AAs) were identified with I_rel_ > 0.1% (0.14-0.35%). In contrast, the most abundant cluster 4 peptides in e.g. 3% Neut (0.44-0.63%) were substantially shorter (12-16 AAs) and only covering the N-terminal part of the γ36 target sequence. This adds to the peptide-level evidence, substantiating why Tryp, but also Alc, PPHs have significantly higher emulsifying activity than Neut PPHs.

Peptide-centric analysis further indicates that mapping identified peptides onto a target cluster consensus sequence alone is not enough to describe high surface activity, but that a higher degree of peptide-level detail is needed. It also calls for further investigations to determine which parts of the target cluster sequences constitute the utmost important (core) region for functionality, thereby facilitating efficient peptide mapping that correlates with observed functionality. This may be addressed computationally through development of more sophisticated predictors, able to identify not only core regions but also other structural features of importance for emulsifying activity in addition to amphiphilicity, such as hydrophobic anchors. Moreover, estimating the abundance/concentration of a specific peptide merely by the MS1 intensity is a very rough estimate, as intrinsic and sequence specific properties makes peptides behave and ionize differently in MS (Jafarpour, Gregersen, et al., 2020; Jarnuczak et al., 2016; Sinitcyn, Rudolph, & Cox, 2018). This highlights the need for fundamentally new computational approaches for absolute peptide quantification in highly complex mixtures such as hydrolysates, which is currently under investigation in our lab. This would allow a more accurate evaluation of peptide release and the direct, quantitative comparison between peptides rather than using raw MS1 estimation, which may be biased on the single peptide-level.

Interestingly, high intensity cluster 1 and 3 Alc peptides (Fig. 5 and Table A.3) indicate that Alc has strong preference to cleave after particularly Glu and Leu in both the N- and C-terminal. Similar trends were observed for high intensity cluster 4 Alc peptides (Table A.4). While Leu specificity was described by the manufacturer, Glu specificity was not. Nevertheless, this is in line with previous reports on Alc specificity (Doucet et al., 2003; Lu et al., 2021). These inconsistencies highlight a crucial aspect for successful application of the presented methodology. In addition to the need for core region mapping and accurate peptide abundance estimation, a high degree of insight on protease specificity is pivotal for making accurate *in silico* analysis and prediction of peptide release. This task is significantly easier to perform for highly specific proteases (e.g. trypsin), also highlighting the need for further development of high specificity industrial proteases, available in bulk amounts for cost-effective process design.

### 3.6. Protease and E/S ratio governs differential protein family selectivity in heat denatured PPI

Using our previously published method for relative protein abundance estimation using length-normalized, relative MS1 intensity (Gregersen et al., 2021, 2022), it is possible to obtain insight on which proteins are abundantly represented in a hydrolysate and thereby also on protein-level enrichment and selectivity. By summing intensities for all peptides originating from the proteins, the potential peptide-level bias is alleviated to a large degree. This is the prerequire assumption used in the iBAQ approach for relative protein quantification (Schwanhüusser et al., 2011). A similar approach was recently developed for non-conventional bottom-up proteomics data, where a specific protease has not been applied in the sample preparation and downstream data analysis (Gregersen et al., 2021, 2022), thereby making the approach suitable for the MS data obtained in this study. Based on relative abundance by I_L_^rel^, protein-level abundances were pooled into major protein families/classes found in potato (Fig. 6, Table A.5). Compared to a previous study on the same PPI (García-Moreno, Gregersen, et al., 2020), patatin is enriched in Alc PPHs while somewhat depleted in Tryp PPHs. In fact, patatin represents more than double of the relative protein content in Alc PPHs (40-45%) compared to Tryp PPHs (18-19%) at 1% and 3% E/S ratio. The direct opposite is observed for Kunitz peptides, where Tryp PPHs at 1% and 3% E/S ratio contain twice the relative amount (46-54%) compared to Alc PPHs (23-25%). As both Alc and Tryp are serine endoproteases (Peyronel & Cantera, 1995), this is likely a direct result of different specificities, and that Alc is capable of hydrolysing patatin to a much higher degree prior to inhibition by Kunitz serine protease inhibitor (KTI-B class) activity remaining despite heat treatment. This observation also correlates well with why the relative protein class distribution for Neut PPHs to a much higher extent reflects earlier MS-based proteomics studies of the PPI (García-Moreno, Gregersen, et al., 2020), as Neut is a Zinc-protease and limited inhibitory activity is expected in the PPI, as discussed in Section 3.2.

**Fig 6:**
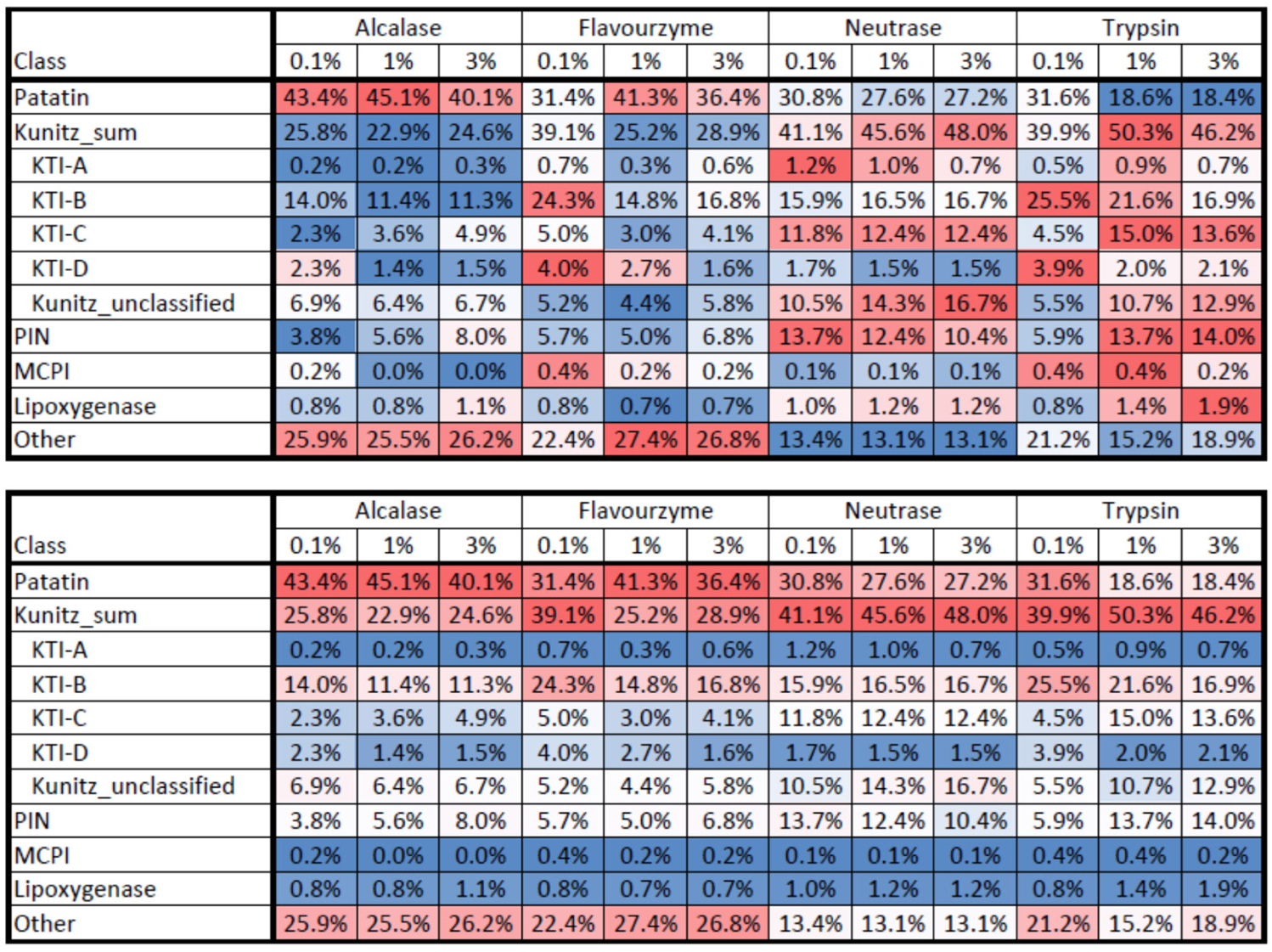
Heat map (blue (low) to red (high)) of relative protein abundance (by unspecific I_L_^rel^) according to protein families/classes for all PPHs (freeze-dried supernatant after hydrolysis). Heat map color is normalized by row (top) and column (bottom) for inter- and intra-sample comparison, respectively. All indented Kunitz subclasses (A-D and unclassified) are included in the “Kunitz_sum” abundance, but are listed explicitly to distinguish quantitatively between subclasses.

For all subclasses of Kunitz-type inhibitors and the class proteinase inhibitors (PIN), substantial differences are observed across PPHs and in comparison to our previous study on native PPI. This is of particular interest for the patatins, KTI-A, and KTI-B classes, as these represent all the target peptides (Table 1). As ten of the 15 target peptides originate from patatin isoforms, enrichment of patatin-derived peptides may be a direct reason for the strong emulsifying properties of the Alc PPH (Fig. 6), even though Tryp PPH was both predicted to have better emulsifying properties as well as shown to contain peptides with better target peptide overlap in the high abundance target clusters 1 and 3. This also illustrates that there is room for improving the interfacial properties of Tryp PPHs even more, by improving hydrolysis conditions and obtaining a higher relative amount of patatin-derived peptides in the PPH. This may potentially be accomplished by combing enzymatic hydrolysis with other methods such as e.g. ultrasound and microwave treatment, previously shown to improve digestibility of potato protein (Cheng et al., 2017; Falade, Mu, & Zhang, 2021; Mao, Wu, Zhang, Ma, & Cheng, 2020).

In the class of “other” protein, which are substantially overrepresented in Alc and Flav PPHs (Fig. 6), the most abundantly quantified proteins (I ^rel^ > 1% in at least one PPH) are related to stress response and glycolysis/carbohydrate metabolism (Table A.5). These include, for instance, two induced stolon tip (IST) proteins (P33191 and M1AFN6), which are particularly abundant in the Alc PPHs (6.3-7.7%). Interestingly, the two IST proteins represented <0.005% of the total protein in the same PPI (García-Moreno, Gregersen, et al., 2020), where the PPI was characterized by means of conventional bottom-up proteomics using tryptic in-gel digestion. The two IST proteins constitute <0.7% of the protein in the Tryp PPHs, however only one of 516 peptides identified for P33191 and two of 316 peptides identified for M1AFN6 were fully tryptic (and very low intensity). Consequently, their identification is hence ascribed to the chymotrypsin activity in rTrypsin/PTN (Nongonierma et al., 2017) as the proteins have a very low frequency of tryptic AAs (Arg/Lys). Chymotrypsin activity is absent in pure, sequencing-grade trypsin used for conventional proteomics, explaining the observed discrepancy. In other studies, the two IST proteins were determined to constitute 0.34% of the total protein content in raw potatoes (i.e. not in a isolate/concentrate) (Krutz et al., 2019). Nevertheless, our observations further substantiate how protease specificity and potential selectivity, search parameters, and hydrolysis conditions significantly affect the peptidome of a hydrolysate in a differential manner, as previously reported for bacterial and seaweed protein (Gregersen et al., 2022). This is particularly relevant for short-term, partial enzymatic hydrolysis.

## 4. Conclusion

In line with increasing focus on green transition and clean label foods, peptides and protein hydrolysates attract significant attention for substituting chemical additives as surface active ingredients in foods. With this work, we present a fundamentally novel approach of data-driven targeted hydrolysis, as an alternative to the conventional trial-and-error methodology. Using prior *in vitro* knowledge of highly potent emulsifier peptides derived from abundant potato proteins, we use *in silico* sequence analysis to hypothesize that Trypsin can release target peptides through hydrolysis and produce a hydrolysate with superior interfacial activity. This was verified to indeed by true though assessment of emulsifying and foaming properties and by benchmarking against the native substrate, the gold standard (sodium caseinate), an enriched patatin fraction, and a range of industrial proteases. In fact, only the application of Trypsin was able to improve both emulsification activity and stability significantly (P<0.05), compared to untreated native substrate. Overall, we found a weak relation between degree of hydrolysis and bulk interfacial activity for the hydrolysates, but DH cannot by itself be used to asses emulsification potential. Using LC-MS/MS analysis, we were able to convert conventional bottom-up proteomics into a non-specific peptidomic analysis, identifying more than 10,000 peptides in each hydrolysate. Using peptide mapping, we show that random overlaps is insufficient for quantitatively describing bulk functionality of hydrolysates, but a deeper, peptide-centric analysis is required. Through this, we show that hydrolysates produced using Trypsin, and to some extent Alcalase, were rich in peptides with much higher amphiphilic potential than the other hydrolysates assayed. Moreover, the 3% tryptic hydrolysate was found to contain predicted peptides, thereby not only validating our novel approach for targeted hydrolysis, but also providing peptide-level evidence to why this particular hydrolysate had the best surface active properties across all hydrolysates investigated. Ultimately, based on modest yields, and that peptides from patatin appear depleted in the hydrolysate, we expect that optimizing process conditions will improve the surface active properties of the tryptic hydrolysate even further. This study further highlights several challenges and bottlenecks related to efficient, large-scale application of the methodology. For instance, a method for accurate and absolute peptide quantification is needed, and better characterization of protease specificity as well as a broader selection of high specificity industrial proteases are prerequisites for further development in this direction. Nevertheless, this study is yet another example of how interdisciplinary research, big data, and computational predictions is gaining headway in food science and can pave the way for more efficient development in the future while simultaneously providing a deeper fundamental understanding of molecular mechanisms and properties related to food ingredient functionality.

## Author contribution

SGE: Conceptualization, Methodology, Formal analysis, Investigation, Validation, Writing – original draft preparation, Writing – review and editing, Visualization, Supervision. AJ: Methodology, Formal analysis, Investigation, Writing – original draft preparation, Writing – review and editing, Visualization. BY: Validation, Writing – review and editing. PJGM: Validation, Writing – review and editing. MGP: Methodology, Formal analysis, Writing – review and editing. DH: Methodology, Formal analysis, Writing – review and editing. CJ: Conceptualization, Writing – review and editing, Funding acquisition, Supervision. MTO: Conceptualization, Methodology, Writing – review and editing, Funding acquisition, Supervision. EBH: Conceptualization, Writing – review and editing, Project administration, Funding acquisition, Supervision.

## Acknowledgements

The authors would like to acknowledge KMC AmbA for supplying the potato protein isolate and Lihme Protein Solutions (Denmark) for supplying purified patatin as a reference. Likewise, the authors would like to acknowledge Arla Foods A/S (Denmark) and Novozymes A/S (Denmark) for providing sodium caseinate and proteolytic enzymes, respectively.

