## Supplementary information and data for "Targeted hydrolysis of native potato protein: A novel route for obtaining hydrolysates with improved interfacial properties"

**Table A.1:** Selection of previously published peptide sequences with *in vitro* functional data for study delimitation and *in silico* sequence analysis as part of the process design for targeted hydrolysis. For each peptide, the annotated name in the reference study (Ref) and the single letter sequence are listed. Based on published data, peptides are evaluated by their ability to reduce oil/water interfacial tension (IFT), the mean droplet size (DS) after emulsification, and the physical stability (PS) of the emulsion and ranked from 1 (high potential) to 3 (low potential) as outlined in section 2.1. For each peptide, an average score was calculated and peptides with scores ≤ 2 were selected for sequence analysis.

| Peptide | Sequence | IFT | DS | PS | Score | Select | Ref.* |
|---|---|---|---|---|---|---|---|
| α1 | AKDIVPFYFEHGPHIFNYSGSIIGPMYDG | 2 | 3 | 2 | 2.3 | | a |
| α2 | HHFVTHTSNGARYEFNLVDGAVATVGDPA | 2 | 2 | 2 | 2.0 | Yes | a |
| α3 | KPVSKDSPETYEEALKRFAKLLS | 2 | 1 | 3 | 2.0 | | a |
| β1 | LRVQENALTGTTTKADDASEANMELLVQV | 2 | 3 | 3 | 2.7 | | a |
| γ1 | GIKGIIPAIILEFLEGQLQEVDNNKDAR | 1 | 1 | 1 | 1.0 | Yes | a |
| γ2 | ANMILLVQVGENLLKKSVSEDNHETYE | 2 | 3 | 3 | 2.7 | | a |
| α10 | KKPVSKDSPETYEEALKRFAKLLSDRKKL | 2 | 2 | 2 | 2.0 | Yes | b |
| α11 | EALKRFAKLLSD | 2 | 3 | 3 | 2.7 | | b |
| α12 | DSPETYEEALKRFAKLLSD | 1 | 1 | 2 | 1.3 | Yes | b |
| α13 | NRPFAAAKDIVPFYFEHGPHIFN | 1 | 3 | 3 | 2.3 | | b |
| α14 | AKDIVPFYFEHGPHIFN | 2 | 3 | 3 | 2.7 | | b |
| α15 | IPATILEFLEGQLQEVDNN | 1 | 1 | 3 | 1.7 | | b |
| α16 | ILEFLEGQLQEVDN | 3 | 1 | 3 | 2.3 | | b |
| α17 | KYDGKYLMQVLQE | 2 | 3 | 2 | 2.3 | | b |
| α18 | KYLMQVLQEKLGE | 2 | 3 | 2 | 2.3 | | b |
| α19 | KYLMQVLQEKL | 2 | 3 | 1 | 2.0 | | b |
| β20 | ELDSRLSYRIISTFWGALGGDVYLGKSPN | 1 | 3 | 2 | 2.0 | | b |
| β21 | ELDSRLSYRIISTFWGALGGDVYL | 1 | 3 | 2 | 2.0 | | b |
| β22 | CPFSSDDQFCLKVGV | 1 | 3 | 1 | 1.7 | | b |
| β23 | FIPLSTNIFEDQLLNIQFNIPT | 1 | 3 | 1 | 1.7 | | b |
| β24 | LNIQFNI | 3 | 3 | 3 | 3.0 | | b |
| β26 | GKELDPRLSYRI | 3 | 3 | 3 | 3.0 | | b |
| β27 | LNIQFNIPTPKLC | 1 | 2 | 2 | 1.7 | Yes | b |
| β28 | VHQNGKRRLALVKDNPLDVSFK | 2 | 3 | 3 | 2.7 | | b |
| β29 | IGSSSHFGPHIFEGELLNIQFDIS | 1 | 3 | 2 | 2.0 | | b |
| β30 | DDNFCAKVGVVIQ | 3 | 3 | 2 | 2.7 | | b |
| β31 | LGGDVYLGKSPNSDAPCP | 2 | 2 | 3 | 2.3 | | b |
| γ1 | GIKGIIPAIILEFLEGQLQEVDNNKDAR | 1 | 1 | 1 | 1.0 | Yes | b |
| γ34 | CRDDNFCAKVGVVI | 3 | 3 | 2 | 2.7 | | b |
| γ35 | RDDNFCAKVGVVI | 2 | 3 | 2 | 2.3 | | b |
| γ36 | FDVIGGTSTGGLLTAMITTPNENNRP | 1 | 1 | 2 | 1.3 | Yes | b |
| γ37 | LLTAMITTPNENNRP | 3 | 1 | 3 | 2.3 | | b |
| γ38 | FCLKVGVVHQNGKRRLALVKDNP | 2 | 3 | 1 | 2.0 | | b |
| γ39 | HQNGKRRLALV | 2 | 3 | 3 | 2.7 | | b |
| γ40 | SSDDQFCLKVGVV | 2 | 1 | 1 | 1.3 | Yes | b |
| γ41 | KDNPETYEEALKRFAKLLS | 1 | 3 | 3 | 2.3 | | b |
| γ42 | DTNGKELNPNSSYRIISIGRGALGGDVYL | 2 | 2 | 3 | 2.3 | | b |
| γ43 | NPNSSYRIISI | 3 | 3 | 3 | 3.0 | | b |
| γ44 | DNFCAKVGVVIQNGKRR | 2 | 2 | 3 | 2.3 | | b |
| γ45 | VGVVIQNGKRR | 3 | 3 | 3 | 3.0 | | b |
| γ46 | FAKLLSDRKKLRANK | 3 | 3 | 3 | 3.0 | | b |
| γ47 | TPNENNRPFAAAKDIV | 3 | 3 | 3 | 3.0 | | b |
| γ48 | GIIPATILEFLEGQLQEVDNN | 1 | 1 | 2 | 1.3 | Yes | b |
| γ49 | FCLKVGVIHQNGKRRLALVK | 2 | 2 | 2 | 2.0 | Yes | b |

| | | | | | | | |
|---|---|---|---|---|---|---|---|
| γ104 | GIIPATILEFLEGQLQEVDNNKDAR | 1 | 1 | 1 | 1.0 | Yes | c |
| γ105 | GIIPGTILEFLEGQLQK | 1 | 1 | 1 | 1.0 | Yes | c |
| α106 | SVSEDNHETYEVALKR | 3 | 3 | 3 | 3.0 | | c |
| α107 | DNPETYEEALKR | 3 | 3 | 3 | 3.0 | | c |
| α10 | KKPVSKDSPETYEEALKRFAKLLSDRKKL | 2 | 1 | 1 | 1.3 | Yes | d |
| α12 | DSPETYEEALKRFAKLLSD | 2 | 1 | 1 | 1.3 | Yes | d |
| β22 | CPFSSDDQFCLKVGV | 2 | 2 | 2 | 2.0 | Yes | d |
| β27 | LNIQFNIPTPKLC | 1 | 2 | 1 | 1.3 | Yes | d |
| γ1 | GIKGIIPAIILEFLEGQLQEVDNNKDAR | 1 | 1 | 1 | 1.0 | Yes | d |
| γ75 | GIIPAIILEFLEGQLQEVDNNKDAR | 2 | 1 | 1 | 1.3 | Yes | d |
| γ76 | GIIPAIILEFLEGQLQEVDNNK | 2 | 1 | 1 | 1.3 | Yes | d |
| γ36 | FDVIGGTSTGGLLTAMITTPNENNRP | 2 | 1 | 1 | 1.3 | Yes | d |
| γ38 | FCLKVGVVHQNGKRRLALVKDNP | 2 | 1 | 1 | 1.3 | Yes | d |
| γ40 | SSDDQFCLKVGVV | 1 | 1 | 1 | 1.0 | Yes | d |

* References: a (García-Moreno, Jacobsen, et al., 2020), b (García-Moreno, Gregersen, et al., 2020), c (Yesiltas et al., 2021), and d (García-Moreno et al., 2021).

**Table A.2**: Color parameter of potato protein hydrolysate powder after enzymatic hydrolyzation by different commercial protease enzymes at different E/S ratios, compared with potato protein isolate (PPI), sodium caseinate (SC) and patatin.

| Enzyme type | Enzyme concentration | Color parameters | | |
|---|---|---|---|---|
| | | *L** | *a** | *b** |
| Neutrase | 0.1 | $69.60\pm0.50^{de}$ | $0.55\pm0.05^{gh}$ | $9.93\pm0.10^{h}$ |
| | 1.0 | $72.77\pm0.47^{d}$ | $0.76\pm0.15^{efg}$ | $10.66\pm0.17^{g}$ |
| | 3.0 | $70.28\pm0.27^{c}$ | $1.30\pm0.08^{d}$ | $11.20\pm0.07^{f}$ |
| Alcalase | 0.1 | $72.39\pm0.79^{d}$ | $0.87\pm0.23^{ef}$ | $11.13\pm0.21^{f}$ |
| | 1.0 | $73.93\pm0.36^{c}$ | $0.54\pm0.09^{gh}$ | $11.48\pm0.25^{ef}$ |
| | 3.0 | $74.31\pm0.61^{c}$ | $0.97\pm0.20^{e}$ | $12.66\pm0.28^{c}$ |
| Flavourzyme | 0.1 | $68.17\pm0.88^{f}$ | $1.64\pm0.18^{c}$ | $11.23\pm0.13^{f}$ |
| | 1.0 | $65.01\pm1.50^{g}$ | $2.16\pm0.09^{b}$ | $11.86\pm0.40^{de}$ |
| | 3.0 | $61.18\pm0.42^{h}$ | $3.05\pm0.17^{a}$ | $12.06\pm0.28^{d}$ |
| Trypsin | 0.1 | $68.62\pm0.44^{ef}$ | $1.63\pm0.12^{c}$ | $11.28\pm0.04^{f}$ |
| | 1.0 | $70.00\pm0.28^{c}$ | $0.70\pm0.06^{fg}$ | $9.98\pm0.22^{h}$ |
| | 3.0 | $71.83\pm0.58^{d}$ | $0.39\pm0.02^{h}$ | $10.06\pm0.25^{h}$ |
| PPI* | | $85.93\pm0.19^{b}$ | $-2.51\pm0.11^{j}$ | $14.29\pm0.14^{b}$ |
| SC** | | $98.70\pm0.15^{a}$ | $-6.03\pm0.03^{k}$ | $11.80\pm0.04^{de}$ |
| Patatin | | $86.63\pm0.19^{b}$ | $-2.74\pm0.05^{i}$ | $15.50\pm0.21^{a}$ |

*Potato protein isolate (native, before heat treatment)
**Sodium caseinate
Mean±SD. all data are based on three replicates
Different small superscript letters in each column indicate the significant differences among means at 95 confidence level (α=0.05)

**Table A.3:** Top 10 highest intensity peptides with >50% overlap with target peptide cluster sequences (cluster 1+3) for all PPH samples along with relative MS1 abundance (by $I_{rel}$) for each peptide. For each PPH, the cumulative, relative abundance of top5 and top10 peptides (by $I_{rel}$) is listed. Table appended as separate .xlsx file.

**Table A.4:** All peptidelevel data from MaxQuant analysis of LC-MS/MS data (peptides.txt). The MaxQuant table (peptides.txt) can also be found in the linked ProteomeExchange repository dataset. Table appended as separate .xlsx file.

**Table A.5:** All protein-level data from MaxQuant analysis of LC-MS/MS data (proteinGroups.txt). The MaxQuant table (proteinGroups.txt) can also be found in the linked ProteomeExchange repository dataset. Table appended as separate .xlsx file.

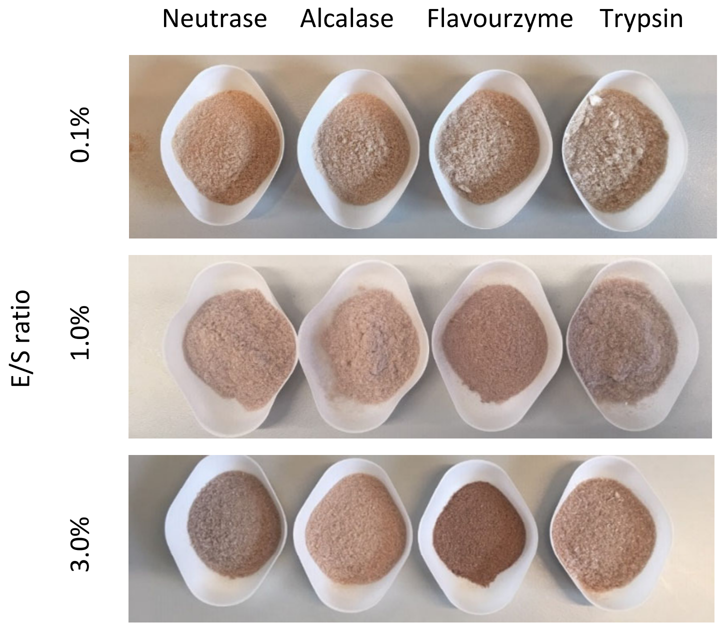


**Fig. A.1:** Visual appearance of freeze-dried PPH from enzymatic hydrolysis of PPI by different proteases (left to right) at different E/S ratios (top to bottom).

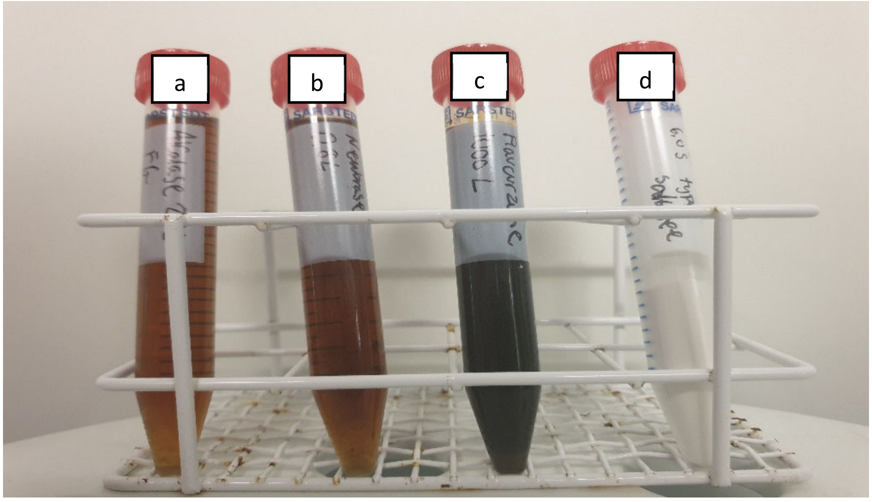


**Fig. A.2:** Visual appearance of enzymes used for proteolysis of PPI: a) Alcalase 2.4 L FG, b) Neutrase 0.8 L, c) Flavourzyme 1000L, d) Pancreatic Trypsin Novo (rTrypsin).

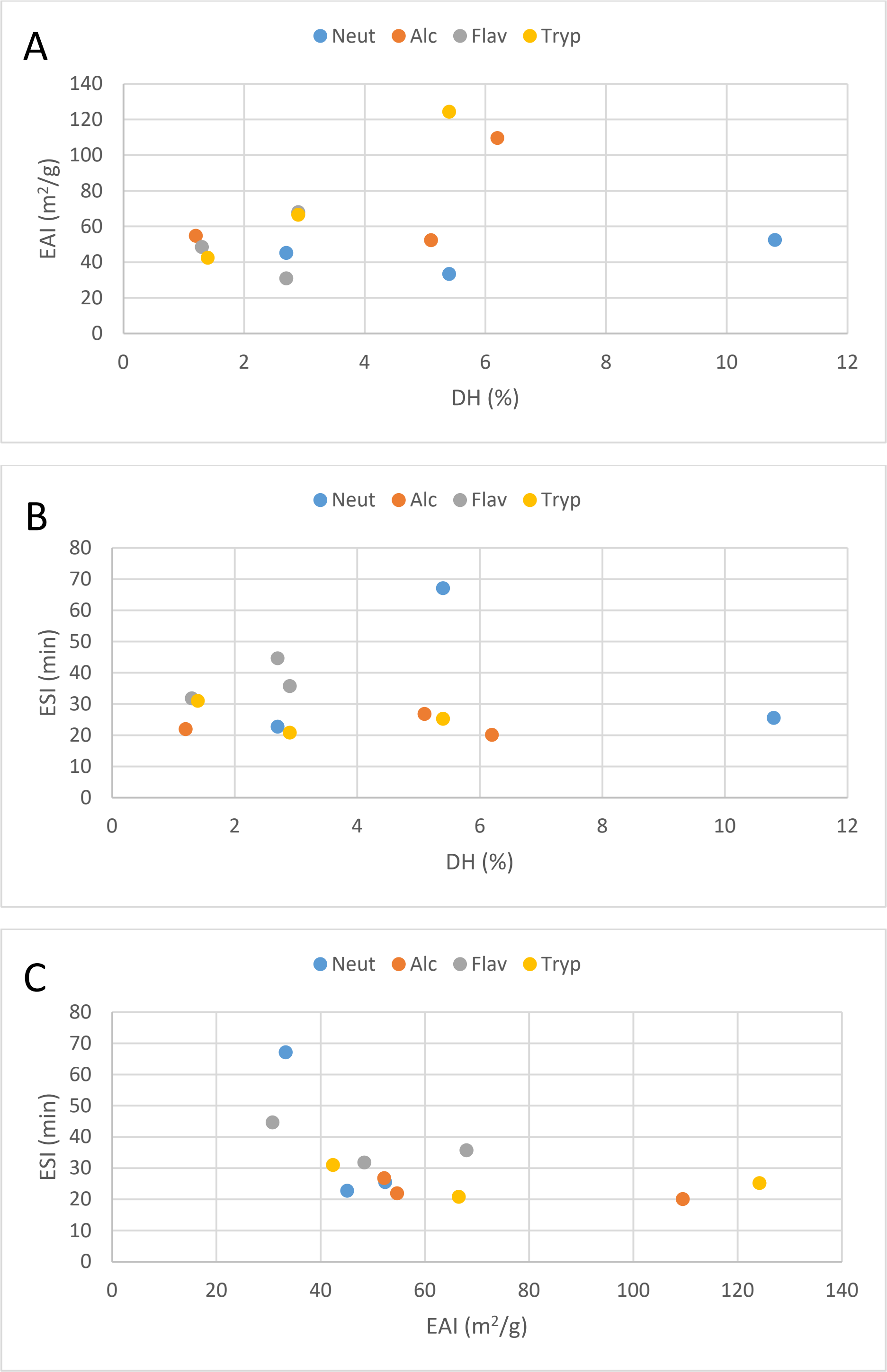


**Fig. A.3:** Scatter plots for investigating potential correlation between DH vs. EAI (A), DH vs. ESI (B), and EAI vs. ESI (C) for PPHs obtained in this study.

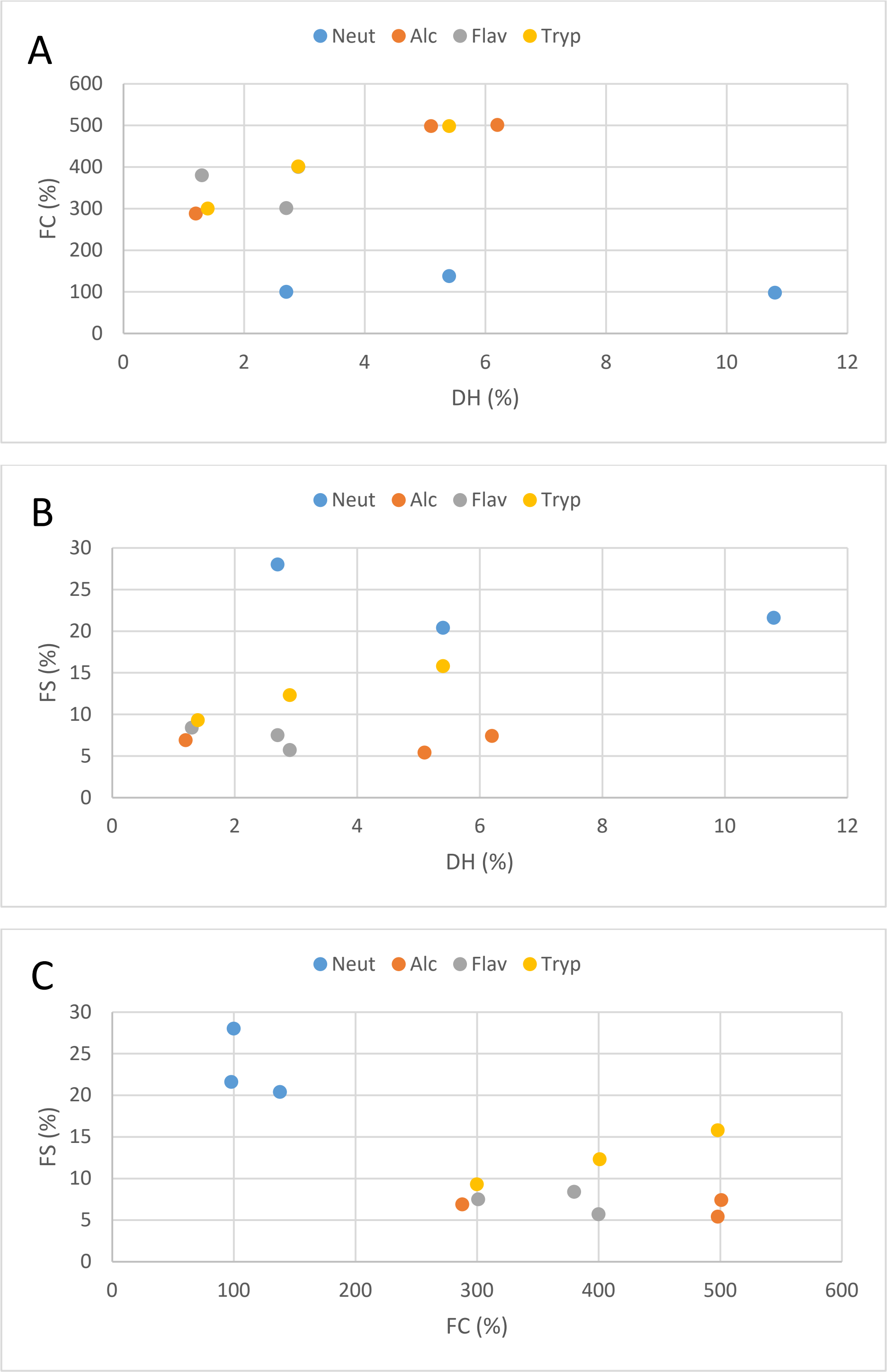


**Fig. A.4:** Scatter plots for investigating potential correlation between DH vs. FC (A), DH vs. FS (B), and FC vs. FS (C) for PPHs obtained in this study.

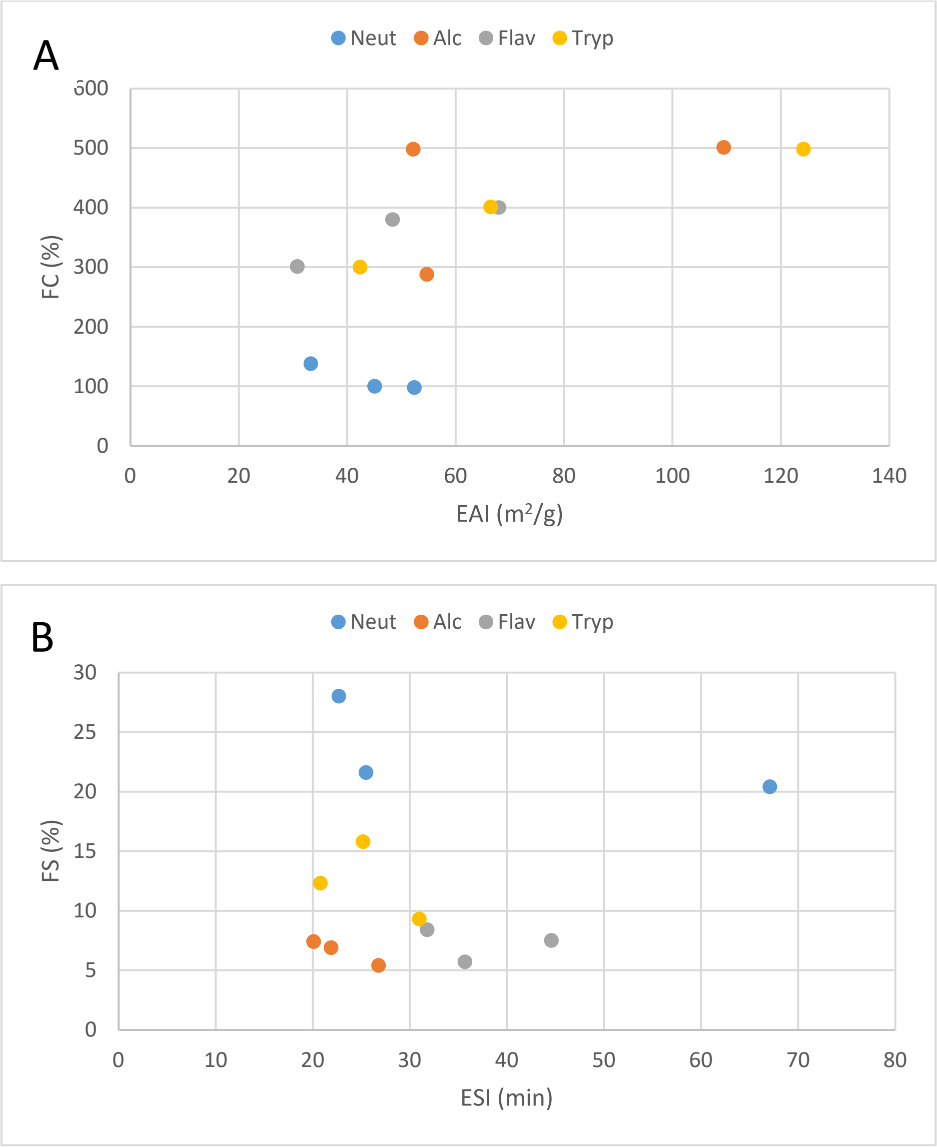


**Fig. A.5:** Scatter plots for investigating potential correlation between EAI vs. FC (A), and ESI vs. FS (B) for PPHs obtained in this study.

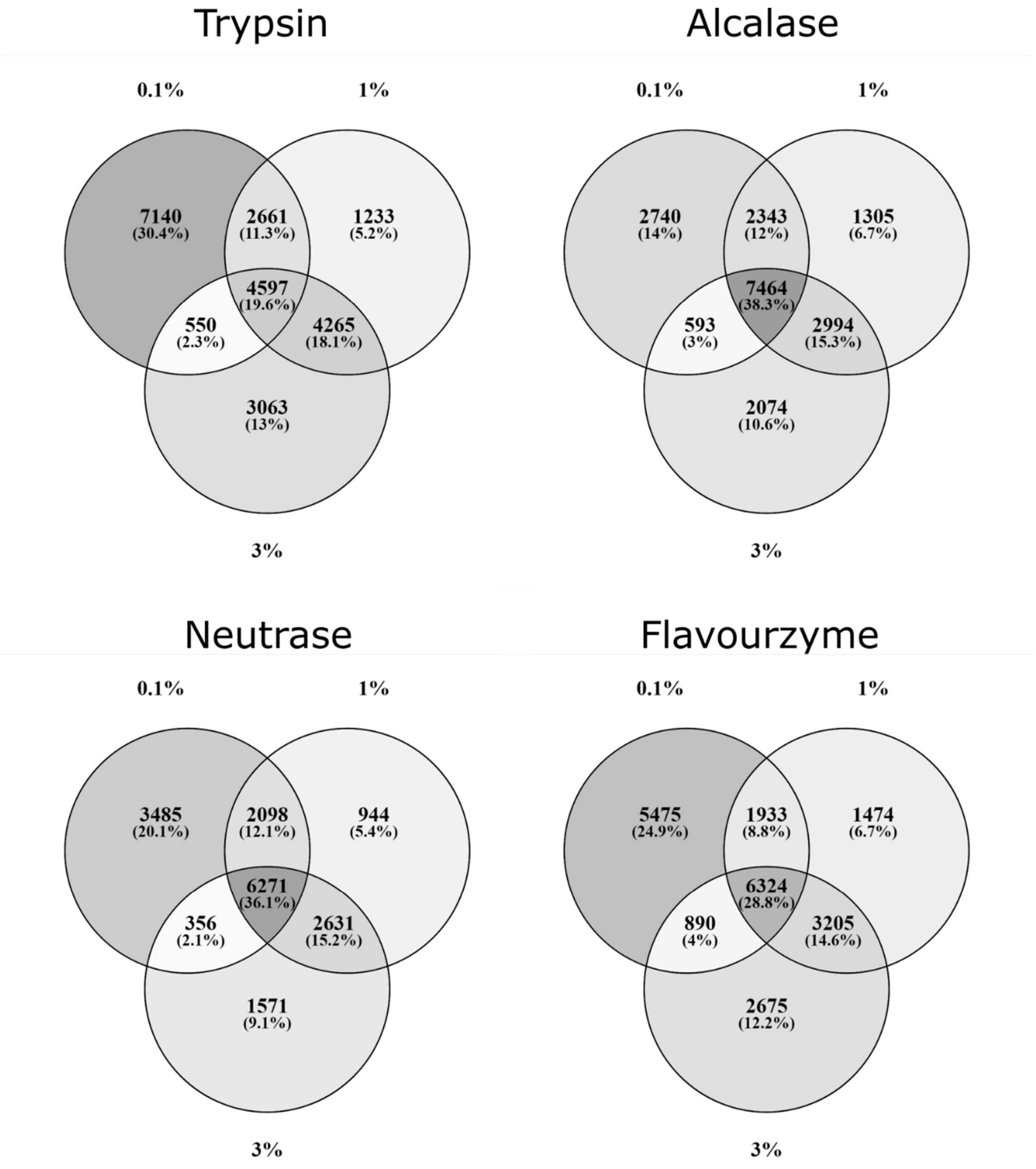


**Fig. A.6:** Venn diagrams showing the number of shared peptide identifications between PPHs obtained using varying E/S ratio (0.1%, 1%, and 3%) for Trypsin, Alcalase, Neutrase, and Flavourzyme.

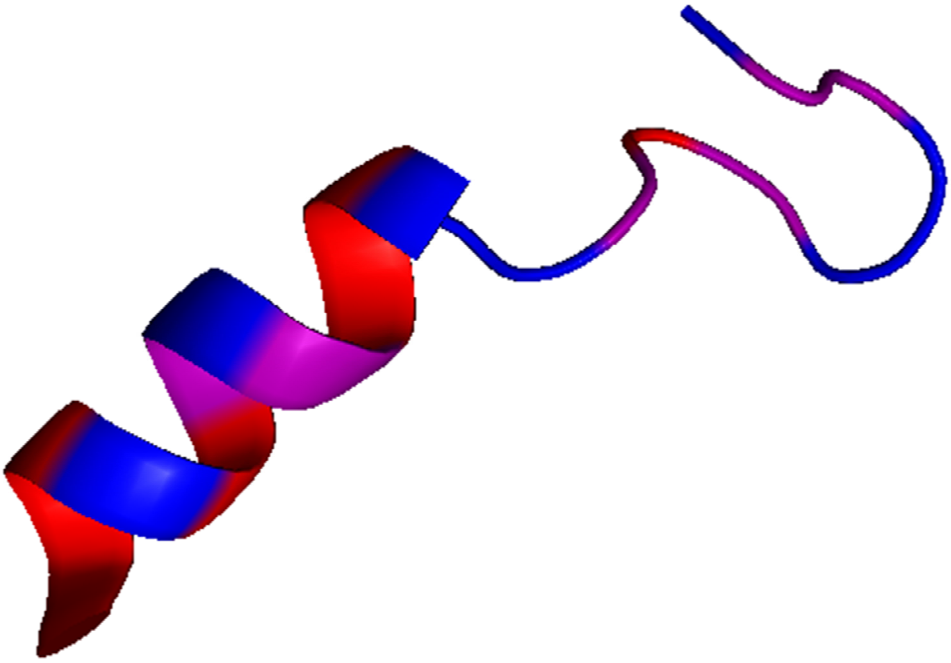

**Fig. A.7:** Native conformation in Patatin-B2 (P15477) of the high abundance Cluster 3 Alc PPH peptides (LLAQVGENLLKKPVSKDNPE, top peptide in Fig. 5) covering the region Leu341-Glu360 (left to right). Peptide is visualized in PyMol 1.5.0 and colored according to the Swiss-Model color scheme: Residues are depicted as very hydrophilic in blue, partially hydrophilic in purple, partially hydrophobic in pink, and very hydrophobic in red.
